# Pathophysiology of Dyt1 dystonia is mediated by spinal cord dysfunction

**DOI:** 10.1101/2022.05.05.490750

**Authors:** Amanda M. Pocratsky, Filipe Nascimento, M. Görkem Özyurt, Ian J. White, Roisin Sullivan, Benjamin J. O’Callaghan, Calvin C. Smith, Sunaina Surana, Marco Beato, Robert M. Brownstone

## Abstract

Dystonia, a neurological disorder defined by abnormal postures and disorganised movements, is considered to be a neural circuit disorder with dysfunction arising within and between multiple brain regions. Given that spinal circuits constitute the final pathway for motor control, we sought to determine their contribution to the movement disorder. Focusing on the most common inherited dystonia, DYT1-*TOR1A*, we confined a conditional knockout of *Tor1a* to the spinal cord and dorsal root ganglia (DRG) and found that these mice recapitulated the phenotype of the human condition, developing early onset generalised torsional dystonia. Physiologically, these mice bore the hallmark features of dystonia: spontaneous contractions at rest, excessive sustained contractions during voluntary movements including co-contractions of motor antagonists, and altered sensory-motor reflexes. Furthermore, spinal locomotor circuits were impaired. Together, these data challenge current understanding of dystonia, and lead to broader insights into spinal cord function and movement disorder pathophysiology.

## Introduction

Neural circuits that control movement are distributed across the neuraxis and are comprised of multiple interconnected loops involving the cerebral cortex^1^, basal ganglia^2,3^, thalamus^4^, cerebellum^5^, brain stem^6^, and spinal cord^7^. While each of these loops has its own function, it is the collaboration of the ensemble that ultimately produces functional movement and hence behaviour. When dysfunction develops within or between these loops, movement disorders arise.

The dystonias comprise the third-most common of the movement disorders (after Parkinson’s disease and essential tremor), and are characterised by involuntary sustained muscle contractions across multiple muscle groups, manifesting as abnormal posture and disorganised movements^8^. The irregular muscle activity leading to these hallmark postures and movements bears 3 main neurophysiological biosignatures: (a) spontaneous muscle contraction at rest^9^; (b) excessive, sustained contractions during voluntary movements often involving co-contractions of antagonistic muscles, which may lead to pain in addition to dysfunctional movement^9,10^; and (c) altered involuntary sensory-motor reflexes^11,12^.

The first link between motor control, movement disorders, and the basal ganglia - a cluster of subcortical nuclei - was drawn in the 1600s^13^. Thereafter, multiple movement disorders were subsequently classified as basal ganglia syndromes throughout the 1800s and early 1900s, including Parkinson’s disease, Huntington’s disease, and dystonia^14–16^. In dystonia, however, limited pathology has been found in the basal ganglia in either humans^17^ or animals^18^. Moreover, there is a significant time delay in the development - and response to treatment - of dystonia when the basal ganglia are involved. Specifically, there is a multi-month lag between acquired injury (such as stroke) of the nuclei and development of dystonia^19^, and a similar lag between deep brain stimulation of the basal ganglia and alleviation of symptoms^20^. Furthermore, not all basal ganglia lesions give rise to dystonia^15^, and attempts to genetically manipulate basal ganglia nuclei to produce mouse models of dystonia have not been overtly successful^21^. To accommodate these findings, dystonia is now commonly considered to be a circuitopathy comprising multiple interconnected brain regions involved in movement, including the basal ganglia, thalamus, cerebellum, and cortex^14^.

Ultimately, all output originating from these regions is mediated via motoneurons in the brainstem and spinal cord that send direct projections to muscles to produce coordinated movements. Spinal motor circuits provide key input to motoneurons; these circuits produce and concatenate the basic syllables of limb movement - syllables that are disorganised in dystonia: muscle contractions across joints, within limbs, and between limbs^22^. Given that the spinal cord is the final common pathway for motor control and that dystonia is defined by its abnormal muscle contractions and movement disorganisation^9^, we sought to determine whether spinal cord dysfunction could be responsible for the pathogenesis of the clinical signs of dystonia.

In this study, we focused on the most prevalent genetic form of dystonia: early onset generalised torsional dystonia, or DYT1-*TOR1A*, which is commonly caused by an in-frame deletion of 3 base pairs in exon 5 of the TorsinA (*TOR1A*) gene^23^. We made a mouse model of DYT1-*TOR1A* dystonia that confines *Tor1a* knockout to spinal cord and dorsal root ganglion neurons. We show that these mice develop functional and physiological signs that mirror those seen in human DYT1-*TOR1A* dystonia, and use these mice to map spinal motor circuit dysfunction. Confining the knockout to dorsal root ganglion neurons does not reproduce the phenotype. We conclude that spinal-restricted knockout of *Tor1a* reproduces the pathophysiology of the human condition. This knowledge opens up a new target - the spinal cord - for the development of strategies aimed at treating DYT1-*Tor1a* dystonia. Our findings also highlight the importance of considering spinal cord circuits in the pathophysiology of movement disorders.

## Results

### Restricting Tor1a deletion to spinal circuits

Unravelling the specific contributions of spinal cord dysfunction to movement disorganisation in Dyt1-*Tor1a* dystonia requires site specificity in *Tor1a* deletion: that is, spinal circuits must be directly affected while supraspinal centres are spared. To this end, we used the established Cdx2::FlpO transgenic mouse model as our genetic entry point to manipulating *Tor1a* in spinal circuits as flippase expression is restricted to the developing spinal cord and dorsal root ganglia (DRG)^24^. Drawing inspiration from a prevailing Tor1a-flox approach^25–27^, we developed a new Tor1a-frt mouse in which exons 3-5 are flanked by frt sites (Fig. 1A; Fig. S1A). Through multigenerational breeding of Tor1a-frt with Cdx2::FlpO mice, we generated a caudal-restricted biallelic “double” conditional knockout (d-cko) of *Tor1a* (Fig. S1B-C). Probing for *Tor1a* and torsinA protein expression in brains, lumbar spinal cords, and DRGs confirmed the site-specificity of this approach. When compared to FlpO-negative littermate controls (Cdx2::wt;Tor1a^frt/frt^), caudal-restricted *Tor1a* d-cko (Cdx2::FlpO;Tor1a^frt/frt^) mice showed normal *Tor1a*-torsinA expression in brain (and heart and liver), but a virtual absence in lumbar spinal cords and DRGs (Fig. 1B-C, Fig. S1D-F), thus validating the site-specificity of our strategy.

**Figure 1.**
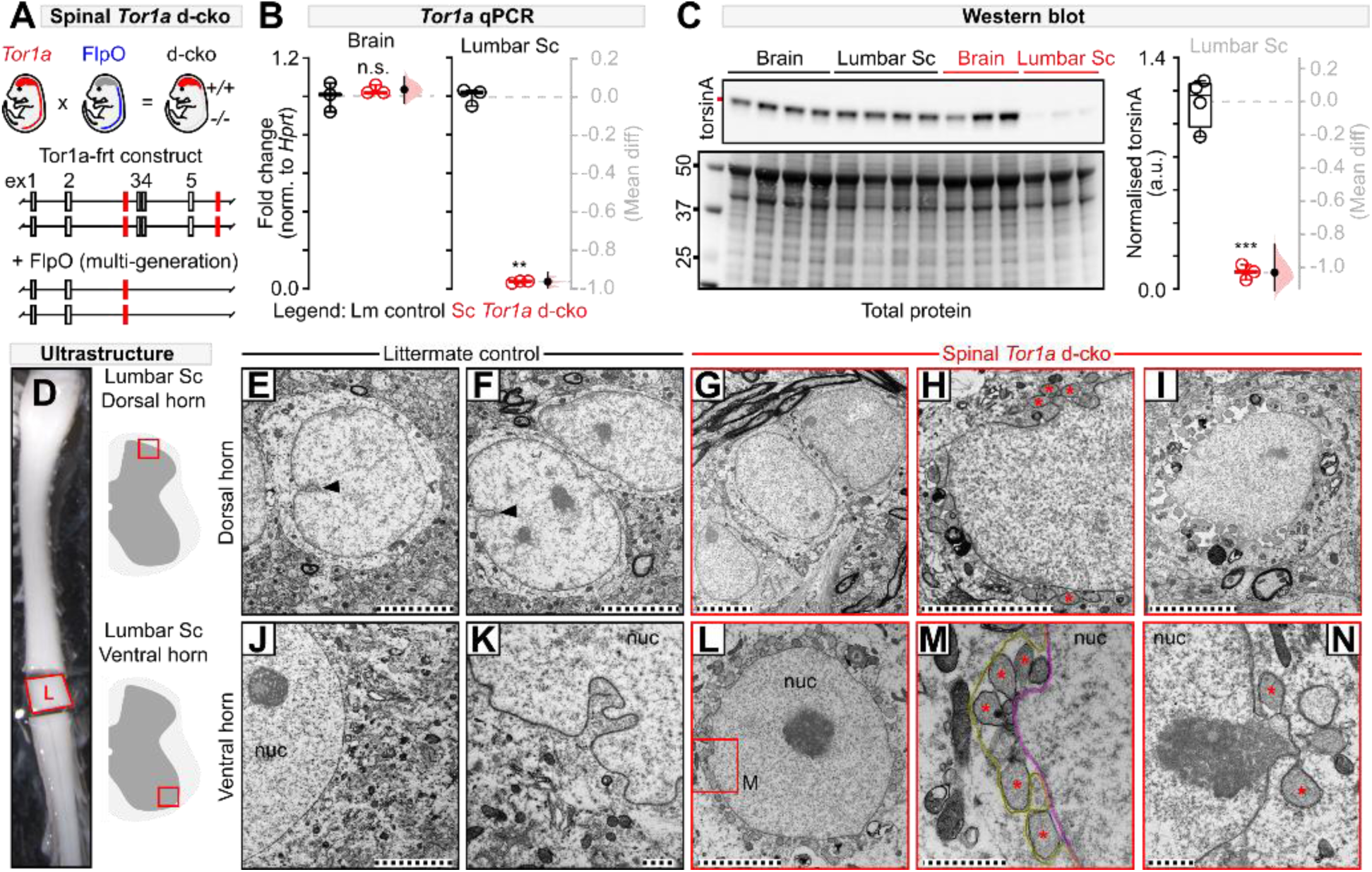
Validation of the spinal-restricted double conditional knockout (d-cko) of *Tor1a*. (**A**) Genetic strategy. (**B**) Unaffected *Tor1a* expression in bains of spinal *Tor1a* d-cko mice (*P*=0.51, independent t-test; *N*=3 P18 control, *N*=3 spinal *Tor1a* d-cko). *Tor1a* expression is absent in lumbar cords of spinal *Tor1a* d-cko mice (\*\**P*<0.001). (**C**) Western blot for torsinA (~37.5 kD) expression in brains and lumbar cords of control (*N*=4) and spinal *Tor1a* d-cko mice (*N*=3). TorsinA expression is abolished in lumbar cords of P18 spinal *Tor1a* d-cko mice (\*\*\**P*<0.0001). (**D**) Lumbar cords isolated for ultrastructural analyses of dorsal (**E**-**I**) and ventral (**J**-**N**) neurons (P18 *N*=4 control, *N*=4 spinal *Tor1a* d-cko mice). (**E**-**F**, **J**-**K**) Control dorsal (**E**-**F**) and ventral (**J**-**K**) neurons show normal nuclear membrane morphology with occasional nuclear invaginations (arrowhead). Dorsal (**G**-**I**) and ventral (**L**-**N**) neurons in spinal *Tor1a* d-cko mice show NE abnormalities, including perinuclear accumulation of vesicles (asterisk) and separation between the inner (**M**, magenta) and outer nuclear membranes (**M**, yellow). “nuc” = nucleus. Scales: 5µm (E-J, L), 1µm (K, M-N). Group data shown (box plots) with individual values overlaid (circles) and mean differences (Gardner-Altman estimation plots). Statistics details in Table S1. See also Figure S1.

### Nuclear envelope pathology in Tor1a-deleted spinal neurons

We next screened for anatomical biosignatures of torsinA dysfunction: nuclear envelope (NE) malformations. Canonical torsinA expression in neurons is distributed throughout the endoplasmic reticulum and NE, but in loss-of-function mutations, torsinA aberrantly accumulates in the NE^28^. Morphologically, this can lead to outer nuclear membrane protrusions that balloon into the perinuclear space where they are released as vesicles^26,29,30^. We found that, while littermate control spinal neurons had normal, well-defined, closely-apposed nuclear bilayers with occasional nucleoplasmic reticulations decorating the nuclei (Fig. 1D-F, J-K, arrowhead), *Tor1a*-deleted lumbar spinal neurons were chockfull of NE abnormalities. In the dorsal horn, there were groups of spinal neurons that appeared normal (Fig. 1G), exhibited early signs of NE budding with sparse vesicle accumulation (Fig. 1H), or showed vesicle-packed perinuclear space with overt separation of the nuclear membranes (Fig. 1I). In the ventral horn, almost all spinal neurons screened were affected, with the perinuclear space filled with NE-derived vesicles (Fig. 1L). Multiple vesicles often budded from one protrusion point of the inner nuclear membrane (Fig. 1M) with signs of electron-dense chromatin content filling the vesicles and being released (Fig. 1N). Large area ultrastructural analysis via backscatter scanning electron microscopy of all contiguous spinal neurons embedded within hemicord slices corroborated the ventro-dorsal gradient in NE severity (available at open access data repository). Of the 2,711 spinal neurons screened, approximately 60% showed ultrastructural abnormalities consistent with torsinA loss of function (Fig. S1W,Y). In contrast, less than 10% of the homonymous DRG neurons screened were affected (Fig. S1H-L, W-X). That is, spinal neurons are particularly vulnerable to *Tor1a* dysfunction. Moreover, ultrastructural screening of basal ganglia neurons - the historical epicentre of dystonia pathogenesis - did not show aberrant NE budding or vesiculation (Fig. S1M-V). Together, these data confirm that this new model confines the *Tor1a* d-cko to spinal and DRG neurons, of which the former manifests overt anatomical biosignatures of torsinA loss-of-function. Henceforth, we will therefore refer to this model as “spinal *Tor1a* d-cko mice.”

### Spinal-restricted Tor1a leads to severe early-onset, generalised dystonia

Severe DYT1-*TOR1A* is defined by a core set of clinical features; most salient are the eponymizing motor symptoms: early onset, generalised spread of disorganised movements^31^. Motor dysfunction emerges early in infancy to childhood, usually at a lower extremity^31^. Over time, motor signs generalise, spreading rostrally to the trunk and upper limbs, stabilizing at or below the neck, usually sparing cranial muscle function^32^. We discovered a striking similarity between severe DYT1-*TOR1A* and spinal *Tor1a* d-cko mice (Fig. 2A-H). The motor impairments emerged early, within the first 1-3 days post-birth, manifesting caudally as hindlimb hyperextension (Fig. 2A; Movie S1). These signs spread bilaterally to affect both lower extremities by postnatal day 5 (Fig. 2B). With increasing age the motor impairments spread rostrally such that by postnatal day 7-9 there were clear signs of pelvis, trunk, and forelimb dysfunction, with the forelimbs abnormally extended forward and minimal body weight support (Fig. 2C-D). By postnatal day 11, the motor signs became fixed at or below the head, sparing orofacial movements. Stepping was impaired as indicated by excessive hindlimb hyperextension with minimal flexion (Fig. 2E). Hindpaw clasping and truncal torsion occurred during tail suspension (Fig. 2F; Movie S2), a test commonly used in Dyt1-*Tor1a* dystonia animal models to uncover latent dystonic-like behaviours^33,34^. By postnatal day 19-21, spinal *Tor1a* d-cko mice were severely dystonic with profound disorganisation of limb movements during voluntary behaviour (Fig. 2G; Movie S3), with interrupting bouts of debilitating truncal torsion (Fig. 2H; Movie S4). Similar to severe DYT1-*TOR1A*, postures were abnormal (Movie S5) and movements were disorganised, jerky, and tremulous in spinal *Tor1a* d-cko mice (Movie S6).

**Figure 2.**
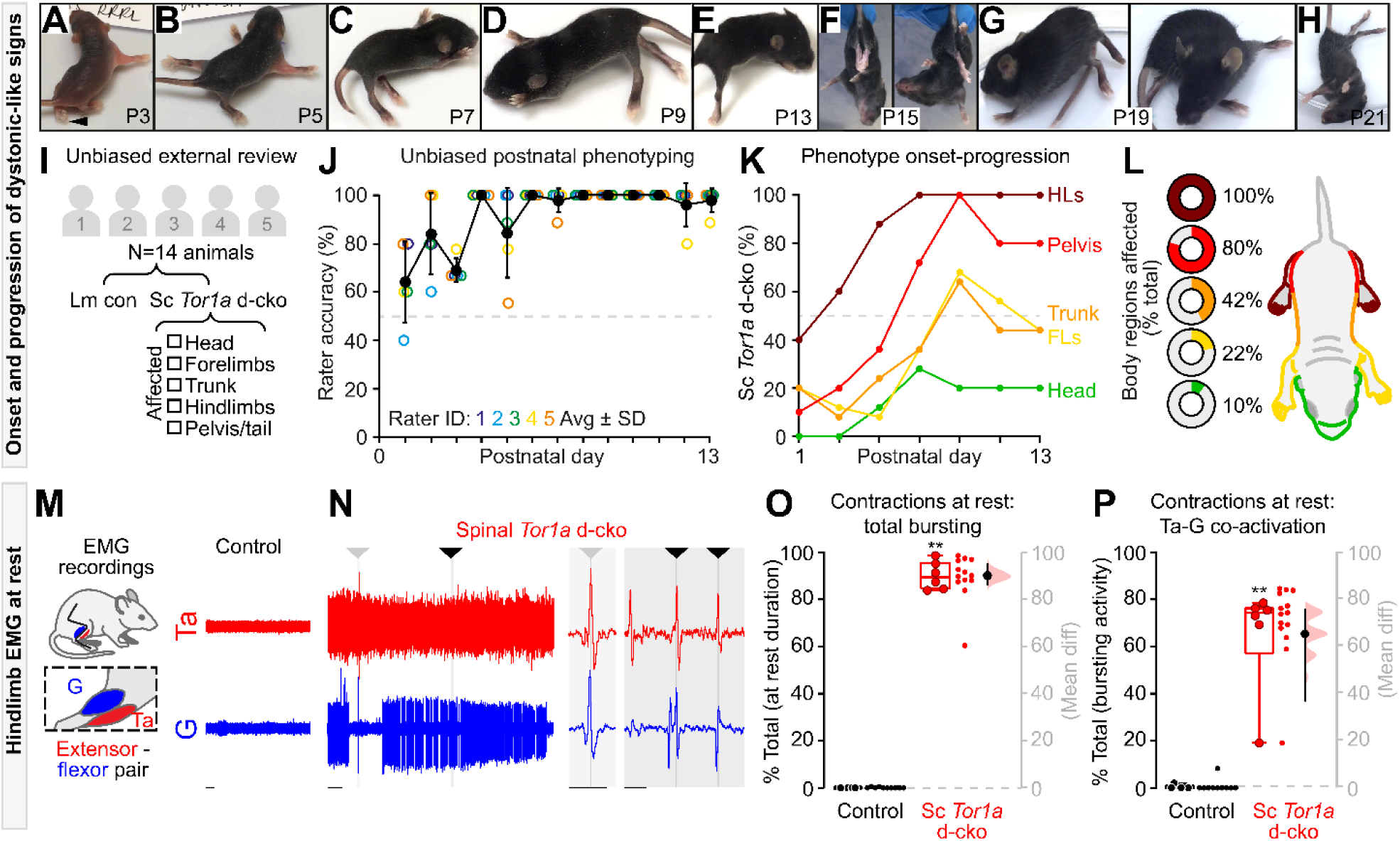
Spinal-restricted *Tor1a* d-cko leads to early-onset, caudo-rostral progression of movement disorganisation marked by abnormal muscle activity. (A-H) Onset-progression of dystonic-like signs in spinal *Tor1a* d-cko mice. (A) Signs emerge early (P1-P3) as hindpaw flexion issues (arrowhead) then (B) spread rostrally and bilaterally during postnatal maturation leading to hindlimb hyperextension, and (C-D) generalise to affect trunk and forelimbs. Affected body regions show (E) increased tone, (F) hindpaw clasping and trunk torsion, (G) abnormal posturing, and (H) truncal torsion. (I) Experimental design for unbiased external peer review of phenotype (*N=*8 control, *N=*6 spinal *Tor1a* d-cko mice). (J) Rater accuracy. (K-L) Spatiotemporal progression of the spinal *Tor1a* d-cko phenotype. (M-P) EMG recordings from gastrocnemius (G) and tibialis anterior (Ta) (*N*=6 P17 and P19 spinal *Tor1a* d-cko mice). (M) Representative EMG activity during rest from P18 wildtype control mouse. (N) Spinal *Tor1a* d-cko mice show excessive spontaneous EMG activity at rest, including coincident single unit spikes (arrowheads, expanded view in shaded regions). (O) Quantification of spontaneous contractions observed at rest (N=4 Control, N=6 spinal *Tor1a* d-cko mice). (P) Quantification of Ta-G co-contractions during the at rest spontaneous EMG activity. \*\**P*<0.01, \*\**P*<0.01, Mann-Whitney U test. Group data shown (box plots) with individual values overlaid (circles) and mean differences (Gardner-Altman estimation plots). Dots: all at-rest epochs analysed/animal. Scale: 1s (M-N), 0.1s (N, shaded regions). Statistics details in Table S1. See also Figure S2 and Movies 1-7.

Given that Cdx2::FlpO directs recombination to both spinal and DRG neurons^24^, we set out to determine whether *Tor1a* dysfunction confined to DRGs alone (so-called DRG *Tor1a* d-cko mice) contributes to early onset generalised dystonia. Leveraging an analogous multi-generational breeding strategy, we confined the biallelic knockout of *Tor1a* exons 3-5 to DRG neurons using the established Advillin-cre^35^ and Tor1a-flox models^34^, respectively. After confirming that *Tor1a* was deleted from DRGs and spared in the spinal cord, postnatal video recordings were performed. In stark contrast to the spinal *Tor1a* d-cko model, DRG *Tor1a* d-cko mice did not develop early onset generalised torsional dystonia (Movie S7, Table S1). That is, *Tor1a* dysfunction in the spinal cord leads to the phenotype of early onset, generalised dystonia in spinal *Tor1a* d-cko mice.

After establishing that spinal-restricted *Tor1a* d-cko causes an early onset, generalised movement disorder, we set out to unambiguously define the spatiotemporal window of the dystonic-like phenotype. Five external raters experienced with mouse behaviour were recruited to provide unbiased analyses of postnatal sensory-motor development in littermate controls vs spinal *Tor1a* d-cko mice (Fig. 2I), all while blinded to the study design, disease model, mutation, and anticipated motor impairments. Raters assessed postnatal video recordings (postnatal day 1-13) and selected - via a unidirectional online test - whether the mouse was a “control” or “mutant,” the latter prompting a follow-up question to select the body region(s) affected. The unbiased external analysis corroborated our internal findings. At postnatal day 1, the accuracy rate (proportion of correct observations) in the unbiased detection of spinal *Tor1a* d-cko mice was >60% (Fig. 2J). Throughout postnatal maturation, the accuracy rate steadily increased until it reached 100% at postnatal day 7 with any subsequent inaccuracy due to false positives (P1-P6: 73% sensitivity, 88% specificity; P7-P13: 100% sensitivity, 98% specificity). A clear spatiotemporal pattern emerged from these unbiased assessments, with motor impairments noted early in the hindlimbs, then spreading rostrally to affect the pelvis, trunk, and forelimbs, with the head minimally affected (Fig. 2K-L). Thus, *Tor1a* dysfunction confined to spinal circuits causes an overt movement disorder that recapitulates the spatiotemporal motor symptoms of severe early-onset, generalised dystonia.

The disordered postures and movements observed in severe DYT1-*TOR1A* are defined by two biosignatures within the underlying muscle activity: (a) persistent involuntary electromyogram (EMG) activity at rest^9^ and (b) disorganised muscle activity during voluntary movements, often including antagonistic co-contractions^9,10^. To this end, we performed acute EMG recordings from the antagonistic tibialis anterior (Ta) and gastrocnemius (G) hindlimb muscles in pre-weaned wildtype control and spinal *Tor1a* d-cko mice (Fig. 2M, left panel).

At rest, there was little to no evidence of spontaneous muscle activity in control mice (Fig. 2M, right panel). Conversely - and similar to severe DYT1-*TOR1A* - spinal *Tor1a* d-cko mice show excessive spontaneous activity at rest, including prolonged co-active bursting in antagonist muscles (Fig. 2N; Fig. S2B-C) with few periods of quiescence (Fig. S2A). Indeed, over 90% of the total resting EMG activity was marked by hindlimb muscle activity (Fig. 2O), of which 65% was flexor-extensor co-activation (Fig. 2P) - including co-active single units (Fig. 2N, shaded regions). We also noted episodes of whole hindlimb stiffness associated with co-contractions of ankle extensor-flexor muscles (Fig. S2D-K) and a proximal tremulous-like phenotype (Movie S6), a phenomenon also reported in dystonia patients^10^.

Although we were unable to obtain EMG recordings in control neonatal mice given their high level of activity, we could leverage the limited mobility of spinal *Tor1a* d-cko mice to assess EMG activity during tail suspension - a common litmus test for abnormal body posturing in dystonic rodents^33,34^ (Fig. S2L) - and volitional locomotion. Tail suspension uncovered a spectrum of EMG patterns ranging from alternation between flexor-extensor bursting to large amplitude co-contractions, and burst disorganisation between Ta-G that was interspersed with tonic activity and rhythmic co-contractions (Fig. S2M-Q). During volitional stepping, spinal *Tor1a* d-cko mice showed hindlimb muscle activity with multiple bouts of antagonistic co-contractions (Fig. S2R-S). In total, Ta-G co-contractions accounted for over one-third of the bursting activity observed during stepping (Fig. S2U). This is clearly an underestimate of the amount of co-activity: when considering single unit coincident activity in addition to the bursts, the proportion of co-contraction time increased (Fig. S2V-W). Disorganised hindlimb muscle activity was further underscored by a considerable range in frequency and duration of bursts (Fig. S2X-Y). Together, these data reveal that spinal-restricted *Tor1a* dysfunction directly leads to an early onset, generalised dystonic-like movement disorder defined by persistent spontaneous muscle activity at rest and excessive co-contractions during rest and voluntary movements.

### Excessive, disorganised motor output in spinal *Tor1a*-deleted mice

The similarity between severe DYT1-*TOR1A* and spinal *Tor1a* d-cko mice raised a fundamental question: are *Tor1a*-deleted spinal circuits the principal source of excessive spontaneous activity and disorganised motor output? Given that the lower extremities are a primary site for disease onset in severe DYT1-*TOR1A* and spinal *Tor1a* d-cko mice, we performed targeted recordings of neural activity intrinsic to the lumbar enlargement - the neural hub for hindlimb motor control^7^. Lumbar spinal cords were isolated from postnatal day 1-5 mice, thus eliminating the influence of descending systems on spinal motor output and framing the previously-defined window of emerging hindlimb dysfunction. Extracellular electrodes were attached to the caudal and rostral lumbar ventral roots to record electroneurogram (ENG) activity from extensor- and flexor-related spinal motor pools, respectively, during “rest” and during drug-induced fictive locomotion^36^ (Fig. 3A).

**Figure 3.**
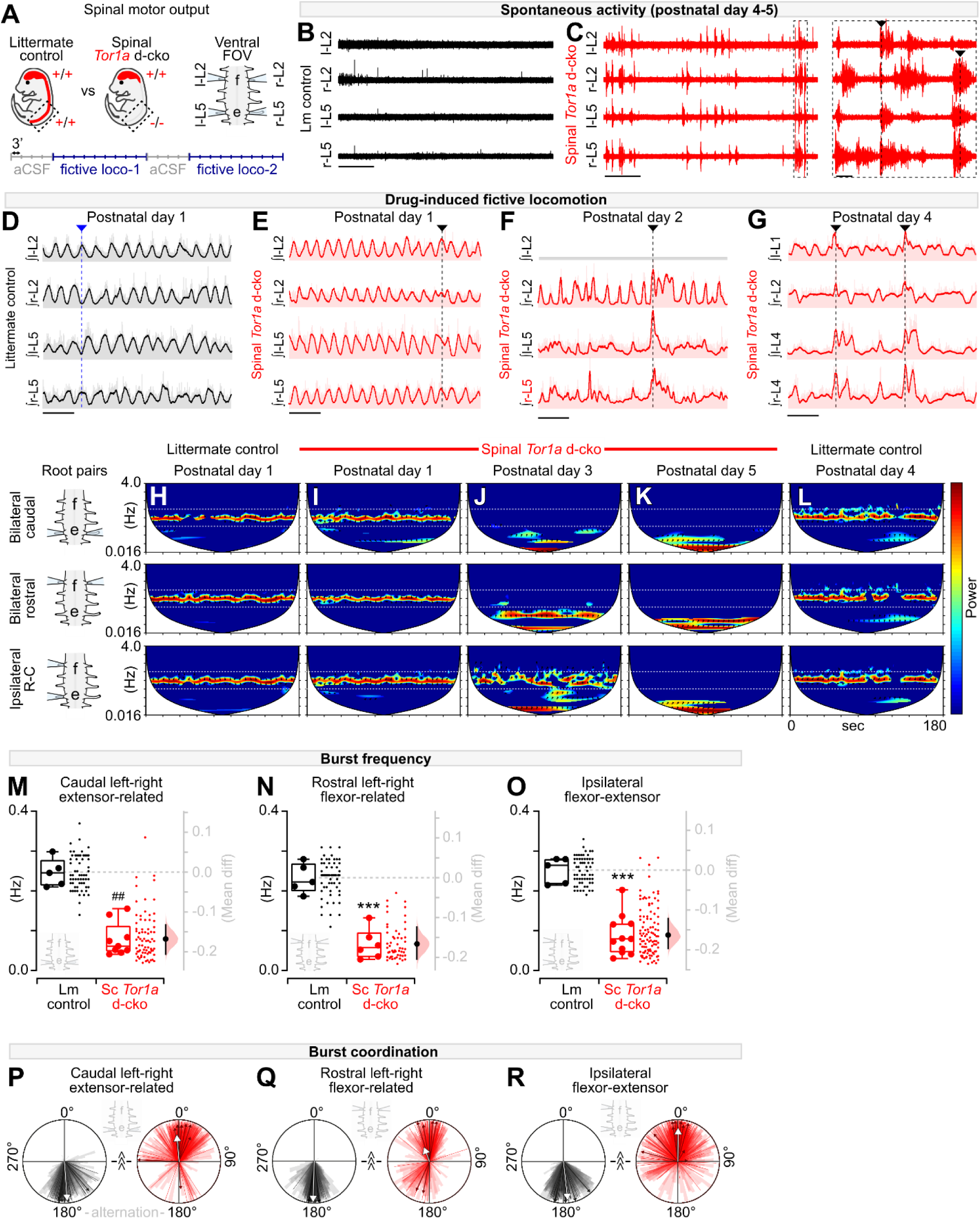
*Tor1a*-deleted spinal circuits produce excessive spontaneous activity and disorganised motor output. (**A**) Experimental design (P1-P5 recordings). (**B-C**) Spontaneous activity at “rest” in artificial cerebrospinal fluid (aCSF). (**D**) Rhythmic, coordinated bursting during drug-induced fictive locomotion in control. Dashed line: alternating bursts at L2, L5, and L2-L5. (**E-G**) Drug-induced fictive locomotion in spinal *Tor1a* d-cko mice. Dashed lines: burst discoordination. Nb: three channel recording shown in F. (**H-L**) Cross-wavelet analysis of frequency-power (colour, blue-red: 2^7^-2^13^) spectrum with phase overlaid (arrows). Horizontal lines: control frequency range. (**M-O**) Burst frequency was decreased in all root pairs assessed in spinal *Tor1a* d-cko (*N*=6-11) vs control (*N*=5) mice (^##^*P*=0.001, Mann-Whitney; \*\*\**P*<0.0001, independent t-test). Group data shown (box plots) with individual values overlaid (circles) and mean differences (Gardner-Altman estimation plots). Dots: all epochs analysed/animal. (**P-R**) Burst coordination was affected in all root pairs (^^^^^*P*<0.001, Watson’s non-parametric U^2^). Bold arrows, orientation: mean phase, length (0-1): concentration of observations. Group data overlaid onto total observations from all epochs (wedges) and epoch averages (lines). Scale: 30s (B-C, left), 10s, (D-G), 1s (C, right). Statistics details in Table S1. See also Figure S3.

In early postnatal isolated spinal cords, there is often spontaneous ENG activity at rest, a transient phenomenon that dissipates until little - if any - unevoked activity is present by postnatal day 4-5^37^, as seen in our littermate controls (Fig. 3B). Spinal *Tor1a* d-cko mice, on the other hand, showed excessive spontaneous ENG activity, including extensive antagonistic co-activation of caudal extensor-related (L5) and rostral flexor-related (L2) motor pools (Fig. 3C-D). These results show that the spontaneous activity recorded in the EMGs *in vivo* could be accounted for by activity in the spinal cord circuits.

Spinal circuits can directly organise and produce rhythmic, coordinated output from flexor-extensor motor pools that manifests as the intra- and interlimb movements defining locomotion^7^. These circuits can be activated *in vitro* by application of neurotransmitters to produce a correlate of *in vivo* locomotion, called fictive locomotion. In littermate controls during fictive locomotion, there is stable rhythmic alternation between bilateral flexor-related (rostral lumbar segments, L2), bilateral extensor-related (caudal lumbar segments, L5), and ipsilateral flexor-extensor related ventral roots (Fig. 3D). In the first 24 hours post-birth, apart from select instances where the normally alternating flexor-extensor activity drifts to synchrony (Fig. 3E, vertical line), spinal motor output in *Tor1a*-deleted mice is largely similar to littermate controls (Fig. S3). But by postnatal day 2, the previously normal alternating flexor-extensor bursting activity becomes disorganised (Fig. 3F; Fig. S3), with prolonged bursting at the caudal extensor-related lumbar motor pools, variable burst durations in the rostral flexor-related motor pools, and co-activation between the flexor- and extensor-related spinal motor pools. By postnatal day 4, ENG bursting was profoundly altered across the lumbar spinal cord (Fig. 3G).

Neural oscillations can be defined by their power, frequency, and phase relationship over time. To determine how the spinal *Tor1a* d-cko fundamentally alters neural output, tiered wavelet transformations were used^38^. We first isolated the dominant power-frequency bands and cycle durations at individual roots (Fig. S3A-E). We then proceeded with a set of cross-root wavelet transformations to extract the shared power, burst frequency, cycle duration, and phase relationships that define: (a) left-right extensor-related (bilateral caudal roots, L4-L5), (b) left-right flexor-related (bilateral rostral, L1-L2), and (c) ipsilateral flexor-extensor related neural activity. The resultant cross-root convolutions were plotted (Fig. 3H-L) and the dominant (high-power) cross-root burst frequency, cycle duration (Fig. S3), and phase relationships over time were extracted for quantitative analysis of spinal motor output.

Littermate controls showed a consistent dominant high-power frequency band confined to 0.125-0.50 Hz for each root pair assessed (Fig. 3H, L). This power-frequency profile was also observed in spinal *Tor1a* d-cko mice at postnatal day 1 (Fig. 3I). But by postnatal day 3 (Fig. 3J), spinal *Tor1a* d-cko mice showed a clear disruption to the signal power-frequency spectrum with a downward shift in burst frequency such that by postnatal day 5, the dominant power-frequency band was ~0.016 Hz (Fig. 3K). Extracting the shared frequencies from the high-power bands revealed a significant decrease in drug-induced burst frequency in all root pairs assessed (Fig. 3M-O). This decrease in burst frequency translated to a ~4-5-fold increase in the cross-root burst cycle duration (Fig. S3F-H).

After establishing the *Tor1a* cko-induced changes to the power-frequency profile, we shifted our focus to cross-root burst coordination, a correlate of the disorganised movements that affect people with dystonia. Cross-root burst coordination data were extracted from the dominant power-frequency bands and plotted on circular graphs wherein 0° denotes in-phase synchrony and 180° reflects out-of-phase alternation.

The classic locomotor profile of out-of-phase bursting activity between bilateral extensors, bilateral flexors, and ipsilateral flexor-extensors^7^ was observed in the littermate controls, with phase data concentrated at 180° (Fig. 3P-R). For the most part, the burst coordination observed in postnatal day 1 spinal *Tor1a* d-cko mice was broadly similar to littermate controls (Fig. S3L). However, this normal bursting profile became disrupted at postnatal day 2-5 wherein there was a predominant shift in the bursting activity towards in-phase synchrony (Fig. 3P-R). Cross-root coherence remained above 0.8 for all root pairs examined (Fig. S3I-K), suggesting that disruption to rhythmic bursting observed in one root was largely related or predictive of the disrupted bursting activity observed in the other root. Together, these data reveal that *Tor1a*-deleted spinal circuits directly produce excessive spontaneous activity at rest and disorganised motor output during locomotion.

### Spinal monosynaptic reflexes are impaired in spinal *Tor1a* conditional knockouts

Given the spinal locomotor circuit dysfunction, we next looked at the most basic spinal circuit, one that can also be readily studied in humans: the monosynaptic (myotatic) reflex. Case reports indicate that individuals with generalised dystonia, including genetically-confirmed DYT1-*TOR1A*, show diminished monosynaptic reflex amplitudes^11,39^, increased variability in the evoked response amplitude^11,12^, and the infiltration of aberrant asynchronous activity^11^. We thus systematically assessed the monosynaptic reflex across space (L1-L5) and time (P7-P13, an age range where the phenotype is fully penetrant) in spinal *Tor1a* d-cko mice.

Graded stimuli were applied to the dorsal root and the evoked monosynaptic reflex was recorded from the ventral root (Fig. 4A, bottom). Plotting representative monosynaptic reflexes revealed a spatiotemporal pattern parallel to the dystonic-like phenotype. Compared to age-matched littermate controls, spinal reflexes in *Tor1a* d-cko mice were reduced in the caudal-most root at postnatal day 7, but otherwise broadly normal in the rostral roots (L1, L3) (Fig. 4B). With increasing postnatal age, the impairments to the monosynaptic spinal reflex spread rostrally, affecting L3 and L1 by postnatal day 11-13. Closer examination of the constituent components of the reflex response (Fig. 4C) revealed that the reflexes in spinal-restricted *Tor1a* d-cko mice had lower amplitudes (Fig. S4F) and longer durations (Fig. 4C-D; Fig. S4G), with multiple asynchronous peaks (Fig. S4A-E). In addition to impairments in the reflex response itself, the latency to onset - a measure largely dependent on afferent conduction - was significantly increased in spinal *Tor1a* d-cko mice compared to littermate controls (Fig. 4G-I; Fig. S4H). Together, these data suggest that there is a clear caudal-to-rostral progression in monosynaptic spinal reflex impairments during postnatal maturation, with a dispersion of the reflex across time.

**Figure 4.**
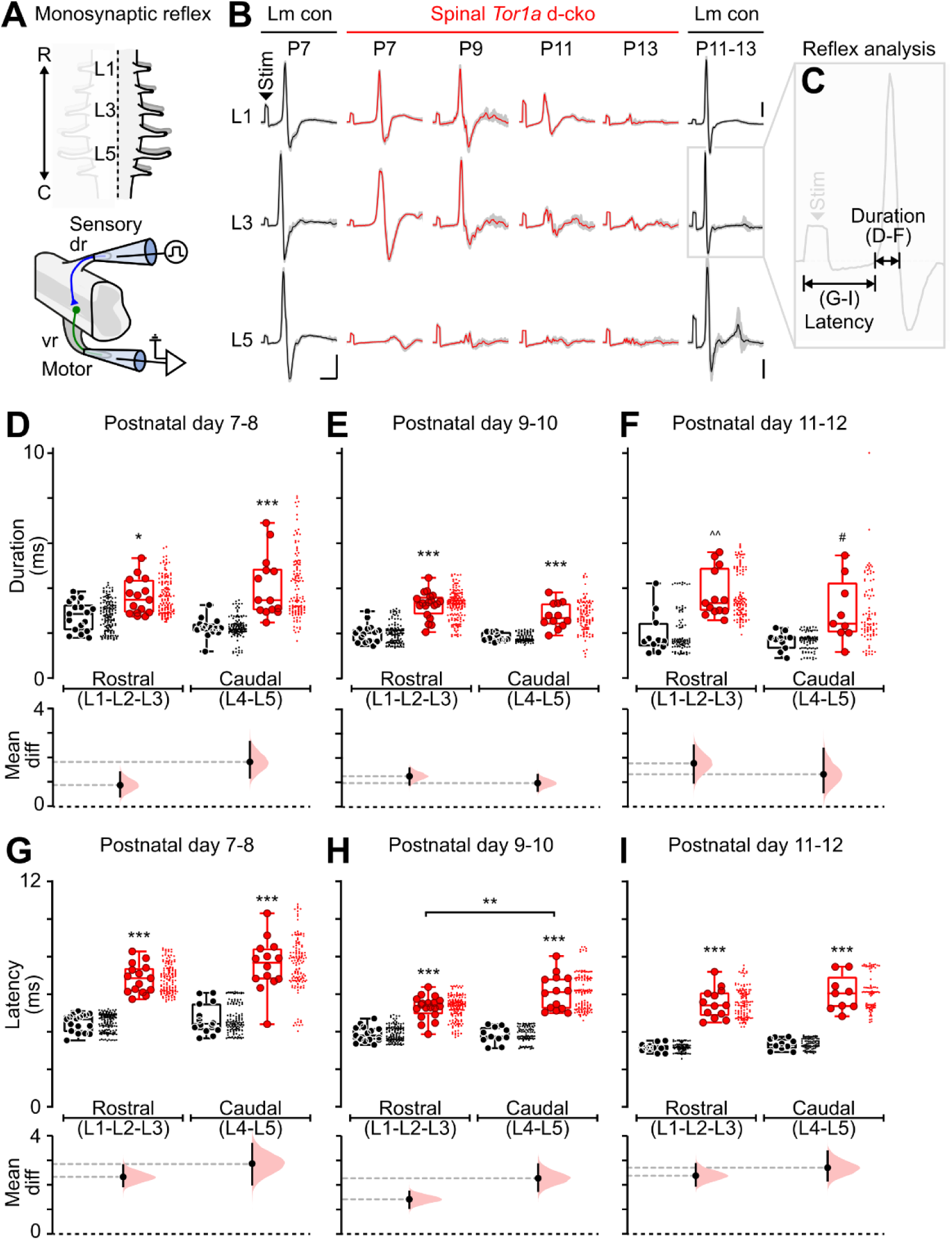
Spinal-restricted *Tor1a* deletion impairs the monosynaptic reflex. (**A**) Experimental design (P7-P13 recordings). (**B**) Representative monosynaptic reflexes. Data shown are average (bold) overlaid onto *N*=10 sweeps (gray). Scale: x=5ms; y=0.05mV. (**C**) Reflex outcome measures. At all ages and roots tested, the monosynaptic (**D**-**F**) response duration and (**G-I**) latency to onset was increased in spinal *Tor1a* d-cko (*N*=21) vs control (*N*=24) mice. \**P*<0.05, \*\**P*<0.01, \*\*\**P*<0.0001, two-way ANOVA and Tukey’s *post hoc* t-test; ^^^^*P*<0.001, Mann-Whitney; ^#^*P*<0.05, independent t-test with Welch’s correction. Group data shown (box plots) with individual values overlaid (circles) and mean differences (Gardner-Altman estimation plots). Dots: all reflexes analysed/animal. Statistics details in Table S1. See also Figure S4.

### Spinal *Tor1a* deletion leads to distributed pathophysiology in the monosynaptic reflex

We next sought to gain mechanistic insights into reflex dysfunction by interrogating the four constituent components of this reflex arc: (a) proprioceptive afferents in the dorsal roots, (b) synapses with motoneurons, (c) the motoneurons themselves, and (d) efferent transmission in the ventral root. We focused on the caudal lumbar motor pools (L4-L5) as they are the earliest affected. We used a ventral horn ablated preparation (Fig. 5A, left) to determine whether motoneurons were intrinsically affected by the spinal-restricted *Tor1a* deletion. Motoneurons in spinal *Tor1a* d-cko mice appeared smaller than those in littermate controls (Fig. 5B). While motoneurons in spinal *Tor1a* d-cko mice had similar resting membrane potentials as control motoneurons (Fig. 5C), there was a significant reduction in whole cell capacitance (Fig. 5D) and a ~350% increase in input resistance (Fig. 5E), consistent with the smaller cell size. Together, these data indicate that lumbar motoneurons are directly affected by *Tor1a* deletion.

**Figure 5.**
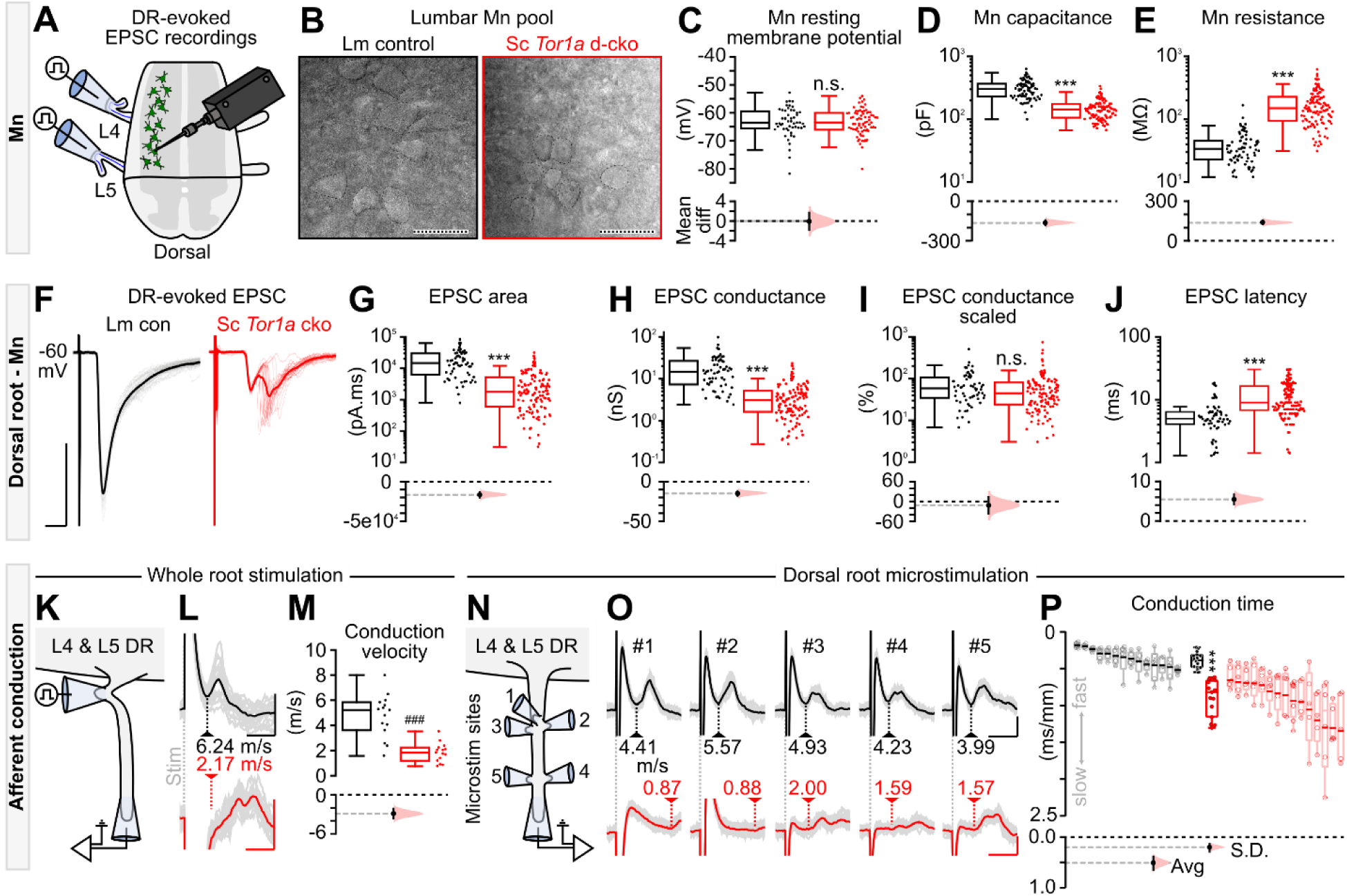
All components of the monosynaptic reflex are impaired in spinal *Tor1a* d-cko mice. (**A**) Experimental design to record afferent- and efferent-evoked excitatory post-synaptic potentials (EPSCs). (**B**) DIC images of P9 lumbar motoneurons (scale: 50µm). For intrinsic motoneuron properties (P1-P13 *N*=# of motoneurons), spinal *Tor1a* d-cko mice showed (**C**) no change in resting membrane potential (*N*=52 vs *N*=77, *P*=0.68, Mann-Whitney), (**D**) decreased whole cell capacitance (*N*=73 vs *N*=108, \*\*\**P*<0.0001), and (**E**) increased input resistance (*N*=73 vs *N*=111). (**F**) P7 dorsal root (DR)-evoked EPSCs. Scale: x=5ms, y=500pA. Spinal *Tor1a* d-cko mice showed (**G**) decreased area (*N*=61 vs *N*=130), (**H**) decreased absolute conductance (*N*=61 vs *N*=128), (**I**) no change in scaled conductance (*N*=61 vs *N*=126, *P*=0.06), and (**J**) increased latency (*N*=60 vs *N*=131) (P1-P13). Group data shown (box plots) with mean differences (estimation plots). Dots: all responses analysed/animal. (**K**) DR stimulation to estimate afferent conduction velocity. (**L**) Representative afferent volleys following whole root stimulation at threshold. Scales: x=0.5ms, y=0.05mV (black) or 0.025mV (red) (L5 DR in P8 control [n=20 sweeps] and P6 spinal *Tor1a* d-cko mice [n=35 sweeps]). (**M**) L4 & L5 DR conduction velocity is decreased in spinal *Tor1a* d-cko mice (P6-P10 *N*=15 vs *N*=15, ^###^*P*<0.0001; independent t-test). (**N-O**) DR microstimulation revealed varied afferent conduction velocities in spinal *Tor1a* d-cko mice (black scale: x=0.5ms, y=0.05mV; red scale: x=3ms, y=0.025mV). (**P**) Afferent conduction time was slower (P6-P10, *N*=14 control vs *N*=15 d-cko mice, Avg: \*\*\**P*<0.0001) and more variable (S.D.: \*\*\**P*<0.0001) in spinal *Tor1a* d-cko mice. Averaged data shown in black and red box plots with individual values (means within each root) overlaid (filled circles) and mean differences (Gardner-Altman estimation plots). The responses from individual roots - including raw (open circles) and averaged (bold line) values - are shown as de-saturated box plots adjacent to the averaged dataset. Statistics details in Table S1. See also Figure S5.

But smaller motoneurons alone could not explain all the changes in the monosynaptic reflex, so we next focussed on the afferent limb of the reflex using low-threshold stimulation of the dorsal roots. Compared to littermate controls, afferent-evoked excitatory post-synaptic currents (EPSCs) in spinal *Tor1a* d-cko mice had reduced amplitude, prolonged duration, and multiple asynchronous peaks (Fig. 5F) - outcomes that were corroborated when we activated a subset of afferent fibres via discrete microstimulations (Fig. S5C-D). At all ages assessed, the EPSC area - a measure of charge carried - was decreased in motoneurons in spinal *Tor1a* d-cko mice compared to controls (Fig. 5G; Fig. S5A). There was a decrease in EPSC conductance (Fig. 5H), but when scaled to input conductance there was no difference (Fig. 5I), suggesting that the monosynaptic effects of afferent inputs to motoneurons are similar for littermate control and spinal *Tor1a* d-cko mice. Of note, at all ages tested, there was a significant increase in the latency to dorsal root-evoked monosynaptic EPSC in spinal *Tor1a* d-cko mice as compared to controls (Fig. 5J; Fig. S5B), a finding that parallels the increased latencies observed in extracellular recordings.

While the longer latencies could result from impairments at the Ia-motoneuron synapse, they could also simply be due to deficits in afferent conduction itself. Thus, we recorded L4 and L5 dorsal root volleys in response to root stimulation (Fig. 5K), and found slower afferent conduction velocities in spinal *Tor1a* d-cko mice compared to controls (Fig. 5L-M), suggesting that the longer latencies to EPSCs resulted from slower conduction velocities. But increased latencies alone cannot account for the asynchronous peaks observed in EPSCs. To this end, we microstimulated the dorsal roots at various sites, activating small subsets of fibres (Fig. 5N-O). After scaling the conduction time by distance, we discovered that spinal *Tor1a* d-cko mice showed longer and variable conduction times as compared to controls (Fig. 5P), suggesting that the multiple peaks in the EPSCs result from time dispersion of the incoming afferent action potentials. That is, two effects occur in the dorsal roots of the spinal *Tor1a* d-cko mice: slower conduction velocities and increased variance of these velocities across fibres.

Given the conduction impairments in dorsal roots, we turned to the ventral roots to determine whether motor axons are also affected (Fig. S5E). We found a modest, but significant decrease in efferent conduction velocity in spinal *Tor1a* d-cko mice as compared to controls (Fig. S5F-G). Responses to microstimulation also revealed increased scaled conduction times and variances in the ventral roots (Fig. S5H-I). In summary, all compartments of the monosynaptic reflex arc - from action potential conduction of sensory afferents to motoneurons to efferent output in the motor roots themselves - are vulnerable to *Tor1a* dysfunction and contribute to impaired sensory-motor integration in *Tor1a* d-cko mice.

## Discussion

### Dysfunctional spinal circuits: a neural substrate for early onset generalised torsional dystonia

We have uncovered that spinal circuit dysfunction is a key contributor to the pathophysiology of DYT1-*TOR1A* dystonia. By confining a *Tor1a* gene mutation to the spinal cord and dorsal root ganglion neurons while leaving expression in the brain normal, mice phenotypically express a generalised torsional dystonia, have an ultrastructural signature indicating loss of function of torsinA^26,29,40^ (with unknown relevance to altered neurotransmission, abnormal muscle contractions, and disorganised movements) in spinal but not brain neurons, have spinal locomotor circuit dysfunction, and have abnormal monosynaptic sensorimotor reflexes.

Co-existing with the motor impairments were signs of sensory dysfunction, with increased variance in the conduction velocities of the fastest dorsal root fibers. While not a major feature of DYT1-*TOR1A* dystonia, sensory abnormalities have been reported on quantitative testing^41,42^, and sensory tricks - or *gestes antagonistes* - can help symptoms of some dystonias^43^. Given that Cdx2::FlpO directs recombinase activity to both spinal and DRG neurons^24^, it is reasonable to ask whether the sensory dysfunction observed in spinal *Tor1a* d-cko mice may be due - in part - to the conditional knockout of *Tor1a* in DRG sensory neurons. That said, few DRG neurons showed nuclear envelope malformations. More importantly, confining the *Tor1a* conditional knockout to DRG sensory neurons does not produce early onset generalised torsional dystonia. Together, these data directly implicate spinal circuits and not primary afferents as the key substrate for dystonia pathogenesis in the spinal *Tor1a* d-cko model.

### The phenotype and pathophysiology of spinal *Tor1a* d-cko mice vs human DYT1-*TOR1A* dystonia

To unambiguously test our hypothesis, we confined a biallelic conditional knockout of *Tor1a* to spinal and DRG neurons, leaving its expression intact throughout the rest of the nervous system. While children with biallelic *TOR1A* mutations are identified with increasing frequency^47–49^, the vast majority of DYT1-*TOR1A* patients have a single allele mutation that is associated with reduced penetrance (~30%) and variable phenotypic expression^50^. Conversely, the spinal *Tor1a* d-cko model shows complete penetrance: 100% of all the genotype-confirmed biallelic knockouts develop early onset generalised torsional dystonia, suggesting that this model may reflect a more fully penetrant form of the human heterozygote condition. Notwithstanding, the spinal *Tor1a* d-cko model is patently similar to the human condition in two key domains: (1) phenotype and (2) physiology.

Phenotypically, motor signs in the spinal *Tor1a* d-cko mice parallel severe DYT1-*TOR1A*: dysfunction emerges early in life in a lower extremity and then spreads in a caudo-rostral fashion during developmental maturation until becoming fixed below the head. Thus far, previous rodent models have not reported or recapitulated this pathognomonic feature of DYT1-*TOR1A*, including conditional-ready models wherein *Tor1a* is manipulated in the cortex^34^, basal forebrain^51^, striatum^27^, or cerebellum^52^ - key nodes in prevailing models of dystonia^14^. Furthermore, in spinal *Tor1a* d-cko mice, the phenotypic signatures of DYT1-*TOR1A* dystonia - abnormal posturing, truncal torsion, and intermittent tremulousness - manifest during naturalistic behaviour when the pups are resting or moving about the environment.

Physiologically, we show that spinal *Tor1a* d-cko mice bear the three primary pathophysiological signatures of DYT1-*TOR1A* dystonia: (a) spontaneous muscle contractions at rest^9^, (b) excessive, sustained contractions during voluntary movements^9,10^, and (c) altered sensory-motor reflexes^11,12^. To date, there has been limited study of the pathological mechanisms underlying these signatures. Equipped with a fully-penetrant model that consistently and reproducibly develops dystonia and a suite of spinal cord preparations to probe sensory-motor dysfunction, we systematically interrogated the precipitating pathophysiological changes of early onset generalised torsional dystonia. Recording from isolated hindlimb motoneuron pools revealed that excessive spontaneous muscle contractions - including co-activation of motor antagonists - can be directly produced by dysfunctional spinal circuits. Much like the *in vivo* phenotype, there are clear caudo-rostral generalisations in spinal circuit dysfunction over postnatal development.

In summary, we identified a dysfunctional neural substrate that phenotypically and physiologically recapitulates early onset generalised torsional dystonia: spinal cord circuits.

### Revisiting evidence in support of spinal circuit dysfunction in DYT1-*TOR1A*

The spinal cord is comprised of neural circuits that control the basic syllables of movement - e.g. reciprocal inhibition to change a joint angle, co-excitation of flexor and extensor motor neurons to stabilise a joint, and co-inhibition of these motoneurons to allow the joint to move freely in biomechanical space^53^. These syllables are concatenated across time to form functional movement^22^. In dystonia, there is abnormal control of these fundamental syllables, akin to a paraphasia of movement. Thus, there was clear logic in considering that spinal circuit dysfunction leads to the signs of DYT1-*TOR1A*.

In fact, there are seeds of substantiation sprinkled through the literature that point to spinal circuit dysfunction. For example, Dyt1-*Tor1a* animal studies have shown nuclear envelope malformations in spinal neurons^26,29^, spinal motoneuron loss^27^, and reduced spinal GABAergic inputs to primary afferent fibres^54^. In non-Dyt1 dystonia models, *Lamb1t* mice have coincident EMG activity between opposing muscles, a phenomenon that persists post-spinal transection and thus directly implicates dysfunctional spinal circuits^55^. And in people affected by DYT1-*TOR1A*, analyses of spinal reflexes indicate that dystonic individuals may have impairments in monosynaptic stretch reflexes^11,12^ and reciprocal inhibition^9,56–58^. Even though these reflexes are mediated by spinal circuits, the impairments observed have been attributed to dysfunction of descending systems^56,59^. But spinal circuits are complex, and form specialised, multi-layered networks that integrate supraspinal, spinal, and sensory inputs to organise motor output^60^. Thus, in dystonia pathophysiology, it is logical to consider spinal circuits as a critical nexus for neurological dysfunction and movement disorganisation in dystonia.

### Reconciling spinal circuit dysfunction with prevailing models of dystonia: movement (dis)organisation at many levels

We have shown that spinal circuit dysfunction can recapitulate one of the most severe forms of primary dystonia. That is, in the homozygous condition, descending command signals cannot override or compensate for spinal circuit dysfunction such that generalised torsional dystonia manifests over postnatal time. Yet one of the most effective treatment options for DYT1-*TOR1A* is deep brain stimulation of a site in the basal ganglia, the globus pallidum interna^61^. If spinal circuit dysfunction leads to disorganised movements, then why is DBS an effective treatment for dystonia? Testing DBS in spinal *Tor1a* d-cko mice is technically unfeasible, due to the combination of rapid onset and progression of motor signs in pre-weaned, undersized pups, the size of the necessary hardware, and the expected duration of stimulation needed for alleviation of dystonic signs. As such, addressing how DBS might be effective if the key pathophysiology is spinal requires revisiting prevailing models of dystonia in the context of motor control at large.

Compared to other DBS-treated movement disorders such as essential tremor or Parkinson’s disease, wherein stimulation offers rapid symptom relief within seconds to hours^62^, many weeks of continuous stimulation is required before tonic dystonic movements show improvement^61^. This delay to symptom amelioration directly implicates neuroplastic mechanisms: a long-term process with adaptive effects that can be localised or distributed via interconnected circuits^62^. In fact, maladaptive neuroplasticity is a widely recognised contributing factor to dystonia^63^ with mis-wired circuitry implicated in the local motor planning ensemble (basal ganglia loops)^64^ as well as distant yet connected circuits such as the corticospinal tract^64^. If dystonia-producing maladaptive plasticity is spatio-temporally distributed across remote yet interconnected circuits (including spinal circuits, as shown here), then dystonia-alleviating adaptive plasticity is likely similarly secured. Indeed, evidence for spinal plasticity has been shown: long-term DBS gradually improved spinal-mediated reciprocal inhibition, restoring agonist-antagonist coordination in the forearms of individuals with generalised dystonia^65^. That is, it seems likely that DBS-mediated improvement of dystonia involves adaptive plasticity throughout interconnected motor ensembles, including spinal circuits.

That spinal cord circuits have the capacity to adapt is not a new thought^66^. Findings over decades of spinal cord injury (SCI) research have established that spinal circuits directly produce organised movements and are intrinsically capable of mediating functional recovery. For example, classic experiments in cats established that following a complete spinal cord transection and resultant paralysis, several weeks of activity-based training can lead to isolated spinal circuits - devoid of descending inputs - to develop the capacity to produce full body weight supported stepping^67–69^. These fundamental studies have been clinically translated: chronic lumbosacral epidural stimulation, paired with activity-based training, can restore function in paralysed humans^70^.

To conclude, with this model of DYT1-*TOR1A* dystonia, we have a newfound entry point into investigating the complex pathophysiology of the disease. As a circuitopathy, dystonia can be considered as a process that affects motor circuits throughout the central nervous system, including those in the spinal cord. The notion that spinal motor circuits are simple relays between the brain and muscles has long been dispelled. Yet spinal circuit dysfunction is rarely considered in movement disorder pathophysiology. We would suggest that new treatment strategies for DYT1-TQRla dystonia could be aimed at addressing the pathophysiology underlying symptoms, the circuits of which are largely resident in the spinal cord.

## Acknowledgements

We are grateful to Nadine Simons-Weidenmaier, Rafaela Fernandez De La Fuente, and Hrista Micheva for assisting in animal husbandry and beta-testing of scoring system; Stuart Martin for initial genotyping and qPCR support; Kim Dougherty (Drexel University), Shayna Singh (Drexel University), Emily Reedich (University of Rhode Island), Cecilia Badenhorst (University of Calgary), and Monica Gorassini (University of Alberta) for serving as our external raters for unbiased phenotyping. We thank Darlene Burke (University of Louisville) for statistical support, Aharon Lev-Tov (Hebrew University of Jerusalem) for providing the SpinalCore software, Martyn Goulding (Salk Institute) for generously providing the Cdx2::FlpO mouse, as well as Tom Warner and Simon Farmer for providing feedback on the manuscript. Thanks to the NVIDIA Corporation for the donation of a Tesla K40 GPU and Quadro RTX 8000.

## Authors contributions

Conceptualization: RMB, AMP

Methodology: RMB, AMP, MGO & FN & MB (patch clamp recordings and conduction velocity testing), CCS & AMP (*in vivo* electrophysiology), IJW (ultrastructure), RS & BJO (qPCR), SS & AMP (Western blots)

Software: AMP, MGO & FN & MB (patch clamp recordings and conduction velocity testing) Validation: AMP

Formal analysis: AMP, MGO & FN (patch clamp recordings and conduction velocity testing)

Investigation: AMP, MGO & FN (patch clamp recordings and conduction velocity testing), IW (ultrastructure), RS & BJO (qPCR), CCS & AMP (*in vivo* electrophysiology), SS & AMP (Western blots)

Resources: RMB, AMP

Data curation: AMP

Writing - original draft: AMP, RMB

Writing - review, editing, approval of final manuscript: RMB, AMP, FN, MGO, MB, CCS, IJW, SS, RS, BJO

Visualization: AMP, MGO & FN (patch clamp recordings and conduction velocity testing) Supervision: RMB

Project administration: RMB

Funding acquisition: RMB (Wellcome, MRC), AMP (EMBO), MB (MRC, BBSRC)

## Declaration of interests

RMB is a co-founder of Sania Therapeutics, Inc and consults for Sania Rx Ltd.

## Inclusion and diversity statement

We support diverse, inclusive, and equitable research.

## Table title and legend

**Table S1. Descriptive statistics.** Con = littermate control, Cdx2::wt;Tor1a^wt/frt^. Cko (red text) = spinal *Tor1a* d-cko, Cdx2::FlpO;Tor1a^frt/frt^. *N=2 outliers (>3 S.D.) removed from dataset prior to analysis (outliers shown in Fig. S4G, L5 dataset). ^#^N=1 outlier removed from dataset prior to analysis (not shown).

## STAR methods

All data generated in this study - including custom written code - are available at our open access UCL Research Data Repository entitled “Pocratsky et al 2022.”

### Animals

All animal procedures were approved by the UCL Animal Welfare and Ethical Review Body and were carried out in accordance with the Animal (Scientific Procedures) Act (Home Office, United Kingdom, 1986) under project license 70/9098 with experiment metadata reported following the NC3R ARRIVE guidelines.

All experiments were performed in pre-weaned male and female mice with date of birth recorded as postnatal day (“P”) 0, or P0. Twenty independent breeding pairs were used to generate all offspring for this study, of which all pups were arbitrarily allocated to the different batch experiments. For longitudinal experiments (e.g. video recordings), pups were randomly selected from the litter for testing. For batch experiments that use the full litter (e.g. electrophysiology), pups were arbitrarily selected from the litter on a day-by-day basis, randomizing the age at which pups were allocated to electrophysiology testing. Experiments were performed while blinded to genotypes, where possible. In the event blinding was impossible at point of data collection (e.g. overt phenotype), data were collected and coded *post hoc* for subsequent blinded analysis. Per experimental design, no spinal *Tor1a* d-cko animals were weaned during this study. Throughout the course of all experiments, the welfare of spinal *Tor1a* d-cko mice was monitored. Spinal *Tor1a* d-cko mice animals could eat (e.g. milk spots), were active, well-groomed, and did not lose weight through the early post-natal period, with most gaining weight (up to ~6g by P19). The most severe phenotype is that shown in Video S6.

Animals were maintained in a vivarium operating under 12-hour light-dark cycle at 19-23°C and 40-60% relative humidity. *In vivo* experiments were performed during the animal’s dark cycle. Animals were group housed (2-4) in individually ventilated cages following the standard husbandry conditions: absorbent bedding (Aspen Chip Eco-Pure Lab), *ad libitum* access to food and water, and environmental enrichment (e.g. nestlets, cardboard tubes, red autoclavable domes, wooden chew sticks and/or balls). For cages housing P14-P21 spinal *Tor1a* d-cko animals, supplemental chow was placed on the cage floor (DietGel 76A with animal protein, ClearH_2_O).

The Cdx2::FlpO strain (Cdx2^tm2(Flpo)Gld^; MGI: 5911680) was generously provided by Professor Martyn Goulding at the Salk Institute (CA, USA)^24^. Upon receipt, FlpO-positive male and female sex chromosomes were transferred into the C57Bl/6J background. The strain was then maintained in the C57Bl/6J background (Charles River) using hemizygous male Cdx2::FlpO mice. The following mice were obtained from the Jackson Laboratory: Advillin-cre (#032536, B6.129P2-Avil^tm2(cre)Fawa^/J; MGI:4459942) and Tor1a-flox (#025832, B6;126-Tor1a^tm3.1Wtd^/J; MGI:5605367). Both lines were maintained in the C57Bl/6J background using heterozygous males after transferring the targeted alleles into the male and female sex chromosomes, respectively.

### Generation of Tor1a-frt mice

The details of the flippase-sensitive Tor1a-frt mouse are summarised in Fig. S1A. Mice were generated by Cyagen Biosciences (CA, USA). To produce the targeting vector, the homology arms and conditional knockout region (*Tor1a* exons 3-5) were generated by PCR using BAC clones RP24-231D2 and RP24-170N21 from the C57Bl/6 library. The targeting vector contained a Neo cassette flanked by loxP sites with DTA used for negative selection. Of the 24 positive embryonic stem cell clones detected with PCR, the 1H5 clone was used to establish the line. Embryonic stem cell injections were performed using C57Bl/6J albino embryos that were then implanted into pseudo-pregnant CD1 females. Founder offspring were identified by their coat colour and germline transmission was subsequently confirmed via breeding with C57Bl/6J females and confirmatory genotyping of the resultant F1 offspring. F1 stocks were transferred to UCL and then bred with C57Bl/6J mice (Charles River) to yield F2 stock with the conditional-ready allele established in males and females. Thereafter, the congenic Tor1a-frt colony was maintained in the C57Bl/6J background using heterozygous Tor1a^wt/frt^ males. Tor1a-frt mice will be made available through the Mutant Mouse Resource and Research Centre (MMRRC).

The following primers were used to detect the pre-Flp conditional-ready allele:

Neo-del-F (F2): 5’-GGCTGGGTTTAGCAGGGAGAAAAG-3’

Neo-del-R (R2): 5’-TAAGACCTGACATGTTTCCTGGGG-3’

The reaction mix contained the following (total 25 µl): 1.5 µl DNA, 1.0 µl of 10 µM forward primer, 1.0 µl of 10 µM reverse primer, 12.5 µl premix Taq polymerase, and 9.0 µl double distilled H_2_O. The cycling conditions were as follows (35 cycles): initial denaturation at 94°C for 3 minutes, denaturation at 94°C for 30 seconds, annealing at 62°C for 35 seconds, extension at 72°C for 35 seconds, and an additional extension step at 72°C for 5 minutes. Genotypes were identified based on the following band sizes (bp): wildtype allele=161, conditional-ready heterozygotes=161 and 306, and conditional-ready homozygotes=308.

### Experimental breeding

Heterozygous Tor1a^wt/frt^ males and Tor1a^wt/frt^ females were bred to produce homozygous Tor1a^frt/frt^ offspring, which were subsequently used to maintain a homozygous line with inbreeding not exceeding 6 filial generations. Homozygous Tor1a^frt/frt^ mice were then used in the following multigenerational breeding strategy to produce offspring with spinal and DRG-restricted biallelic “double” conditional knockout of *Tor1a* exons 3-5 (spinal *Tor1a* d-cko; Fig. S1):

Generation 1 cross: Hemizygous Cdx2::FlpO males bred with homozygous Tor1a^frt/frt^ females to yield spinal *Tor1a* single cko mice wherein exons 3-5 of one *Tor1a* allele are deleted from spinal neurons.

Generation 2 cross: Cdx2::FlpO;Tor1a^wt/frt^ male mice bred with homozygous female Tor1a^frt/frt^ to generate the following offspring used for experiments: (1) Flpo-negative littermate controls (Cdx2::wt;Tor1a^wt/frt^) and (2) spinal *Tor1a* d-cko (Cdx2::FlpO;Tor1a^frt/frt^). Spinal *Tor1a* single cko animals were excluded from this study.

Genotyping of experimental mice (constitutive knockout allele) is as described above with the following primers and product sizes:

Frt-F (F1): 5’-CTAGTGAGGCTCTGGGTAAATGCACAC-3’

Neo-del-F (F2): 5’-GGCTGGGTTTAGCAGGGAGAAAAG-3’

Neo-del-R (R2): 5’-TAAGACCTGACATGTTTCCTGGGG-3’

Band sizes (bp): wildtype allele=161 bp, conditional-ready allele=308, and constitutive knockout allele=216.

An analogous multigenerational breeding strategy was used to generate biallelic *Tor1a* conditional knockout in dorsal root ganglia neurons (DRGs). Specifically, heterozygous Advillin^wt/cre^ mice (vs Cdx2::FlpO) bred with homozygous Tor1a^flox/flox^ mice (exons 3-5 floxed, vs Tor1a^frt/frt^ with exons 3-5 frted). qPCR data validating the site-specificity of the conditional knockout are reported in Table S1.

### qPCR

In total, N=14 P18 mice (7 littermate control 7 spinal *Tor1a* d-cko mice) were deeply anaesthetised via i.p. injection of ketamine:xylazine (300 mg/kg : 30 mg/kg; 0.01 ml/g body weight) and decapitated. Whole brains (with cerebellum attached), lumbar spinal cords, dorsal root ganglia (thoracic-lumbar), hearts, and livers were rapidly harvested, snap frozen in liquid nitrogen, and transferred to −80°C until processing by experimenters blinded to the genotype.

Samples were weighed, digested in a 10% w/v solution of Qiazol lysis buffer (Qiagen RNeasy Lipid Tissue Mini Kit, ThermoFisher 74804), and homogenised using the Ika Homogeniser Workstation with disposable probes. 200 ml of chloroform was added to each sample and mixed vigorously. Samples were then centrifuged at 12,000 G for 15 minutes at 4°C. The top aqueous layer was transferred into a cold falcon tube and an equal volume of chilled 70% ethanol was added. RNA extraction was then performed following manufacturer protocol. A total of 2 µg of total RNA was used for cDNA synthesis (ThermoFisher High-Capacity RNA-to-cDNA kit, 4387406) using the following thermocycler conditions: 37°C (60 minutes), 95°C (5 minutes), and hold at 4°C until plate was stored at −20°C.

Separate TaqMan master mixes (ThermoFisher 4370048) were created for each probe: *Tor1a* (ThermoFisher 4331182, Mm00520052_m1) and reference genes *Actb* (ThermoFisher 4331182, Mm00607939_s1), *Gapdh* (ThermoFisher 4331182, Mm99999915_g1), and *Hprt* (ThermoFisher 4331182, Mm00446968_m1). Samples were run in triplicate using the ThermoFisher qPCR QuantStudio. *Tor1a* fold gene expression values were estimated using ΔΔCt method with *Hprt* or *Actb* serving as the primary housekeeping gene.

The same methods were used for quantifying *Tor1a* expression in the spinal cords and isolated DRGs of N = 6 P58-P59 littermate control (Avil^wt/wt^;Tor1a^wt/flox^) and N = 5 DRG *Tor1a* d-cko mice (Avil^wt/cre^;Tor1a^flox/flox^). Two outliers - one from each group - were excluded from analysis due to potential tissue contamination issues.

#### Western blots

N=7 P18 mice were deeply anaesthetised via i.p. injection of ketamine:xylazine (300 mg/kg:30 mg/kg; 0.01 ml/g body weight) and decapitated. Whole brains (with cerebellum attached) and lumbar spinal cords were rapidly dissected from each animal, snap frozen in liquid nitrogen, and transferred to −80°C until protein extraction. Samples were homogenised on ice in 10% w/v radioimmunoprecipitation assay (RIPA) buffer, then agitated on a rotator at 4°C for 30 minutes. RIPA buffer contained (final concentrations reported): 50 mM Tris-Cl (pH 7), 1 mM EDTA, 2 mM EGTA, 150 mM NaCl, 1% NP-40, 0.5% sodium deoxycholate, 0.1% SDS, and 1x Halt Protease and Phosphatase Inhibitor Cocktail (ThermoFisher 78440). Crude lysates were centrifuged (21,000 G at 4°C for 20 minutes), the supernatant collected, and total protein concentration was estimated using the Pierce BCA Protein Assay Kit (ThermoFisher 23225) with bovine serum albumin as standards. Protein samples were then mixed with 4x Laemmli sample buffer (8% SDS, 40% glycerol, 0.04% Bromophenol blue, 0.25 M Tris HCl [pH 6.8], 10% β-mercaptoethanol), denatured (98°C for 4 minutes), and centrifuged (13,000 G at room temperature [RT] for 5-7 minutes) before electrophoresis. Approximately 20 µg of total protein was separated using NuPAGE 4-12% Bis-Tris protein gel (ThermoFisher NP0321BOX) in MOPS running buffer and then transferred to a 0.2 μm polyvinylidene fluoride (PVDF) membrane using the Trans-Blot® Turbo RTA Mini PVDF Transfer Kit (Bio-Rad 1704272) and a wet transfer approach (Mini Trans-Blot Cell System, Bio-Rad 1658030). After briefly staining with Ponceau S (Sigma P7170-1L) to confirm protein transfer, blots were incubated in blocking buffer (gentle agitation at RT for 60 minutes; blocking solution: 0.1% Tween20 in TBS [“0.1% TBST”] with 7% non-fat dry milk [Marvel Dried Semi-milk <1% fat]) and then probed with rabbit anti-torsinA at 1:1,000 (Abcam ab34540; RRID:AB_2240792) overnight at 4°C with gentle agitation (primary solution: 0.1% TBST, 7% non-fat dry milk, 0.1% sodium azide). The following day, membranes were washed with 0.1% TBST (5 washes at RT, each 5 minutes, gentle agitation), incubated with goat anti-rabbit horseradish peroxidase (HRP)-conjugated species anti-rabbit secondary antibody at 1:1,000 (same solution as primary, RT for 60 minutes, gentle agitation) (Bio-Rad 1706515; RRID: AB_11125142), then washed with 0.1% TBST (6 washes at RT, each 5 minutes, gentle agitation). Immunoreactive bands were detected using HRP substrate (Millipore WBLUC0500) and imaged using the Chemidoc MP Imaging system (Bio-Rad 17001402). Band density was quantified using the Bio-Rad Image Lab software (v6.0.1). Molecular weights were estimated using the Precision Plus Protein Kaleidoscope Standard (Bio-Rad 161-0375). After chemiluminescent imaging, membranes were stained with Coomassie blue R-250 dye to estimate total protein^71^. Using Coomassie blue as a loading control, torsinA blot images were analysed and formatted using Image Lab software.

### Ultrastructure

In total, N=8 P18 mice (4 littermate control 4 spinal *Tor1a* d-cko mice) were deeply anaesthetised via i.p. injection of ketamine:xylazine (300 mg/kg:30 mg/kg; 0.01 ml/g body weight) and transcardially perfused with room temperature 0.1 M PBS (ThermoFisher 70011036) followed by EM-grade 4% formaldehyde (TAAB F003). The skull and vertebral column were harvested and stored in 4% PFA for approximately 3-4 weeks before brains, lumbar spinal cords, and dorsal root ganglia were subsequently dissected.

### Basal ganglia

Whole brains were embedded in 4% agarose (Sigma-Aldrich A4018) and then sectioned coronally using a vibrating microtome (Leica Biosystems VT1200S). Initial sectioning settings of 0.30 mm/s speed, 1 mm amplitude, and 300 µm thickness were used to quickly cut through the brain until the anterior commissure appeared. Thereafter, the cutting thickness was decreased to 200 µm and serial cross-sections were collected until the hippocampal formation became prominent. Slices were stored in a 24-well petri dish filled with chilled 0.1M PBS during collection and then transferred into a 0.1M sodium cacodylate solution containing EM-grade 2% formaldehyde/1.5% glutaraldehyde (TAAB G011) and stored at 4°C overnight. Images of vibratome slices were captured using a stereo microscope to identify which cross-sections contained the globus pallidus and striatum. The globus pallidus and striatum were then bilaterally dissected from two vibratome cross-sections per animal, yielding N=4 globus pallidus and striatum “blocks” per animal, respectively.

Samples were prepared for electron microscopy following a modified protocol^72^. Samples were washed in 0.1M sodium cacodylate buffer and post-fixed in 1% OsO4/1.5% potassium ferricyanide for 60 minutes at 4°C. Samples were then washed in distilled H_2_O and treated with 1% thiocarbohydrazide (TCH) for 20 minutes at RT, 2% osmium (OsO_4_) for 30 minutes at RT, 1% uranyl acetate (UA) overnight at 4°C, and lead aspartate for 30 minutes at 60°C, with intermediate washing in distilled H_2_O between each step. This was followed by dehydration of the samples via increasing concentrations of ethanol: 70%, 90%, and two rounds of 100%. Samples were then infiltrated with 1:1 propylene oxide (PO):epon resin for 60 minutes followed by two rounds of fresh 100% epon for 60-120 minutes before being mounted and polymerised overnight at 60°C. Brain slices were mounted on the flattened top of a pre-polymerised epon stub under a coverslip to maintain the flatness of the tissue/block surface, spinal cord samples were mounted in moulds so that the surface to be sectioned was parallel and adjacent to the final block surface.

### Lumbar spinal cord and dorsal root ganglia

Following dissection from the vertebral column, the L4-L5 lumbar spinal cord segments were identified and the corresponding dorsal root ganglia isolated. Thereafter, each spinal cord was transversely cut to yield N=3, 1 mm blocks (a rostral, central, and caudal block). The rostral and central blocks were treated as described above. All samples were dehydrated via 10-minute incubations of an increasing ethanol gradient: 50%, 70%, 80%, and 100% (x2). Samples were embedded as described above in PO (10 minutes x 3), PO:epon (90 minutes), and then overnight in epon. The following day, samples were placed in fresh epon for 5 hours and then embedded overnight at 60°C. For spinal cord samples, thin slices were then mid-sagittally hemisected to produce left and right hemicords. Each hemicord was small enough to fit onto one grid, allowing the contiguous neurons from dorsal to ventral horns to be visualized. Lower magnification images were captured to confirm the lumbar segment, define the regions of interest for high resolution imaging, and to be used as a reference to map the precise location of each neuron imaged. At the ventral horn, high resolution images were captured at 5 different ROIs with emphasis on the motoneurons. At the dorsal horn, high resolution images were captured at 3 different ROIs along the dorsal-most region.

Samples were trimmed and 70 nm ultrathin sections were cut with a 45° diamond knife (DiatomeDiATOME) using an ultramicrotome (Leica UC7). Sections were collected on 1 x 2 mm Formvar-coated slot grids for transmission electron microscopy (TEM) or on ITO-coated coverslips for scanning electron microscopy (SEM), and imaged using a transmission electron microscope (ThermoFisher Tecnai G2 Spirit) and a charge-coupled device camera (Olympus SIS Morada) or using the backscatter detector on a Gemini 300 SEM (Zeiss), respectively.

SEM imaging was performed using a backscatter detector and at an accelerating voltage of 4.5kV reduced to a landing energy of 1.5kV by application of a stage bias of 3kV, and at a working distance of 6mm. Mosaics were acquired of large areas at 30nm pixel size resolution using Atlas 5 software (fibics).

All TEM and SEM images generated in this study are available at our open access UCL Research Data Repository.

### Quantification

Raw .czi files of spinal cord and DRG SEM micrographs were imported into Bitplane Imaris (v9.7.2) for analysis. All images generated from this study were coded and quantified by an experimenter blinded to the genotype. Using the Spots module, neurons were first identified by a combination of the following ultrastructural features: large euchromatic nucleus, nuclear envelope, perinuclear cytoplasm enriched with subcellular organelles such as Golgi complex and endoplasmic reticulum, and a prominent primary dendrite extending from the cell body. Thereafter, any positively identified neuron was included for analysis if the nucleus and nuclear membrane were visible. Neurons were then further classified as abnormal if one or more of the following morphological signatures were present: nuclear envelope vesiculation, electron dense inclusions, and large putative cytoplasmic inclusions. Due to extensive vesiculation, values reported reflect an underestimate of the number of neurons affected (e.g. nucleus obscured from view due to vesicles). Data shown are the group average and S.D. of percent neurons affected.

### Behaviour

Videos were collected by experimenters blinded to the genotypes from N=4 independent breeding pairs, N=14 mice in total (N=8 littermate control and N=6 spinal *Tor1a* d-cko mice, respectively). In parallel, age-matched video recordings were performed using the control offspring derived from four different dystonia-related strains in the lab (N=16 mice). These videos served as controls for the training dataset for normal postnatal sensory-motor development in mice.

Recordings started at P1-P3 and continued every other day until postnatal day P14-P15, the pre-juvenile stage age at which the primitive reflexes have disappeared and the adult-like sensory-motor behaviours are refined^73^. Prior to weight assessments and video recordings, the parents were transferred to a separate cage while the home cage with pups was placed on an external heating pad. Individual pups were transferred to the clear plexiglass dish of a weight scale and videos were recorded at 30 fps, 1080p using 8 MP camera (f/2.2 aperture, 1.5µm pixel size, 1/3-inch sensor). Approximately 60-90 seconds of video were recorded per pup. Pups were given unique paw tattoos using a hypodermic needle and Ketchum green tattoo ink (F.S.T. 24201-01) following their first recording session to facilitate longitudinal recordings. For each pup, a representative ~30 second segment of video was selected and formatted for external, unbiased peer review.

Five external raters who had previous experience with mouse behaviour were selected to provide unbiased phenotype scoring of littermate control and spinal *Tor1a* d-cko mice.

Raters were informed that they would review postnatal videos of a “new mouse model for a movement disorder” and that they would assign the mouse - via a unidirectional online test - to “control or mutant groups.” If mutant was selected, then raters would “check off which body region(s) are affected.” Thus, raters were blinded to the model (dystonia), the mutation (spinal-restricted conditional knockout), the genotype (littermate vs spinal *Tor1a* d-cko mice), when the phenotype was expected to manifest (postnatal timeline), where signs might emerge along the body axis (e.g. head vs hindlimbs), and how the signs might manifest (e.g. hyperextension, hyperflexion, torsion). Prior to scoring, raters had the option to review a set of annotated training videos that delineated the normal functional milestones typically observed during postnatal maturation in non-disabled pups. The sensitivity of the postnatal testing paradigm was defined as the proportion of spinal *Tor1a* d-cko observations that were accurately classified as “mutant” (N=55 and 115 true positives at P1-P6 and P7-P13, respectively) divided by the total number of true spinal *Tor1a* d-cko observations (P1-P6: N=55 true positives + N=20 false negatives; P7-P13: 115 true positives + N=0 false negatives). The specificity was defined as the proportion of control observations that were correctly classified as “control” (N=101 and N=137 true negatives at P1-P6 and P7-P13, respectively) divided by the total number of control observations (P1-P6: N=101 true negatives + N=14 false positives; P7-P13: N=137 true negatives + N=3 false positives).

A similar recording strategy was used for the DRG *Tor1a* d-cko cohort. In total, N=5 littermate control and N=5 DRG *Tor1a* d-cko mice generated from two independent breeding pairs were recorded from postnatal day 1-9, an age range that reflects the development of stepping as well as the complete onset and progression of early onset generalised dystonia in the spinal *Tor1a* d-cko group. The DRG *Tor1a* d-cko mice did not develop early onset generalised torsional dystonia throughout postnatal maturation and were indistinguishable from littermate controls at P58-P59, the terminal timepoint for fresh tissue collection.

All videos generated throughout the study – including the training dataset, scored videos, and phenotype test - are available at our open access UCL Research Data Repository.

### *In vivo* electrophysiology: hindlimb muscle activity in spinal *Tor1a* d-cko mice

#### Preparation

For each animal, five ~10 cm long pieces of A-M Systems Wire (#793200) were cut to fabricate two sets of bipolar recording electrodes (gastrocnemius and tibialis anterior, respectively) and one monopolar grounding electrode (base of tail). At one end of each wire, approximately 1 mm of coating was removed to expose the underlying wire. At the other end of each wire, approximately ~1-2 mm of wire coating was removed and then inserted into a 30-gauge hypodermic needle extracted from its plastic hub. The needle was then crimped 3-5 times to ensure stable contact between exposed wire and needle shaft. Colour coded heat shrink was attached to the base of the needle to facilitate electrode identification during the subsequent implantation and recording procedure.

The limited size and mobility of spinal *Tor1a* d-cko mice permitted an acute preparation for recording hindlimb EMG activity in pre-weaned mice. N=2 P17 and N=4 P19 spinal *Tor1a* d-cko mice were briefly anaesthetized using isoflurane and ophthalmic ointment was applied to the eyes. The fur overlying the right hindlimb and posterior half of the dorsum was shaved and small incisions were made at the mid-back and at the level of the hip. Underlying adherent tissue between the incision sites was bluntly dissected to create an open channel for routing the recording wires. The 1 mm bared ends of the wires were bent into hook-like shapes using the beveled edge of a 30-gauge needle and then two each were percutaneously implanted into right hindlimb gastrocnemius and tibialis anterior. A monopolar ground electrode was inserted near the base of the tail. The hip incision was loosely sutured closed leaving some excess wire external to the animal to provide sufficient slack for accommodating behavioural activity. After suturing the incisions, the animal was transferred to an open field arena and kept warm with external heating pads until fully alert.

N=4 P18 C57Bl/6J wildtype mice of both sexes were briefly anesthetized with isoflurane until the paw reflex was abolished. While mice were kept warm, the left or right hindlimb was shaved and skin palpated to identify the underlying medial gastrocnemius and tibialis anterior muscles. Bipolar recording electrode wires (A-M Systems Wire #793200) were inserted through the skin and into the belly of the muscle. Anaesthesia was then removed and the mice were transferred to an empty cage enclosed within a custom-made Faraday cage. EMG recordings were performed once the animal recovered from acute anaesthesia. Bouts of stepping and quiet rest were flagged as described below.

#### Recordings

Recordings were made using a custom-made amplifier and headstage. Signals were amplified (100x), bandpass filtered (30 Hz - 10 kHz), and digitized (10-20 kHz) using a CED Power3a running Spike2 v9 software, and saved for offline analysis. Concomitant videos were also recorded for a subset of animals using a Basler camera (acA800-510uc) with a Fujinon lens (DF6HA-1B; 1:1.2/6mm). Videos were recorded at 104 fps, 0.5 megapixel and synched with EMG recordings using manual and software cues interspersed throughout the entire recording session. The duration of recordings ranged from 30-45 minutes and included the following behavioural conditions: recovery from anaesthesia, alert and at-rest, locomotion, and tail suspension. Following terminal recordings, animals were humanely culled and ear biopsies collected for *post hoc* genotyping.

#### Analysis

Using contemporaneous notes, datafile keymarks, and videos as reference, EMG data were segmented into four conditions: at-rest (for baseline EMG thresholding), tremulous-like phenotype at-rest, locomotion, and tail suspension. Noise contamination was removed from the segmented dataset using Spike2 native and custom-written scripts. Contaminants such as glitches (“FixGlitch” script), mains noise (50 Hz notch filter with “HumRemoval” script for 50 Hz harmonics), or motion artefacts (25-50 Hz digital high-pass filter) were eliminated as needed. The resultant clean EMG signals are shown. For locomotor analysis, the cleaned signal was full-wave rectified and subsequently analysed as follows.

For each animal, we first calculated the average at-rest baseline EMG activity level. For subsequent detection of the locomotor-related bursts, we set a minimum rising (burst onset) and falling (offset) threshold crossing to baseline average ± 10 S.D. or average ± 20 S.D., depending on the extent of excessive spontaneous EMG activity at rest. These criteria were visually validated on linear envelope and smoothed EMG signals before proceeding with analysis.

With the established threshold level, the individual EMG events within each burst were detected using the Spike2 “Events” script and then concatenated into full bursts using the “Bursts” script with a maximal burst onset interval of t=0.0025s and longest within-burst inter-spike interval of t=0.010s. From the detected bursts, the following parameters were then extracted for each bout of locomotion (excluding the first and last burst/bout): burst onset-offset, number of events per burst, intra-burst inter-spike interval, intra-burst average inter-spike interval, burst frequency, burst duration, inter-burst interval, and cycle duration. To isolate the high frequency, short duration single unit events from the burst dataset, data were filtered post-event detection to isolate bursts that were <6 events/burst vs ≥6 events/burst. After detecting individual events, which were concatenated into bursts as described above, we then used a custom-written code to quantify the proportion of total bursting activity that was (1) tibialis anterior only, (2) gastrocnemius only, or (3) co-bursting.

Given that the recordings were performed in the open field and spinal *Tor1a* d-cko mice never achieved independent body weight-supported plantar placement, quantification of the classic locomotor measures (e.g. EMG activity related to swing-stance duty cycle) was not feasible. The custom-written code used to process EMG data is available at https://github.com/Brownstone-lab and our open access UCL Research Data Repository.

### *In vitro* electrophysiology: spontaneous activity and drug-induced fictive locomotion

#### Preparation

Fictive locomotion experiments were performed following a modified approach of previously described methods^74^. N=5 P1-P4 littermate controls and N=11 P1-P5 spinal *Tor1a* d-cko pups were decapitated, eviscerated, and the vertebral column isolated from the cadaver. The isolated column was placed in a Sylgard-coated petri dish filled with chilled, continuously bubbled (95% O_2_/5% CO_2_) dissection solution that contained (mMol/L in 18.2 MΩ H_2_O): 215 sucrose (Sigma-Aldrich S1888), 3 potassium gluconate (Sigma-Aldrich P1847-500G), 1.25 sodium phosphate monobasic (Sigma-Aldrich S5011-500G), 26 sodium bicarbonate (Sigma-Aldrich S5761-500G), 4 magnesium sulfate heptahydrate (Sigma-Aldrich 63145), 10 D-glucose (Sigma-Aldrich 158968-1KG), 1 kynurenic acid (Sigma-Aldrich K3375-5g), and 1 calcium chloride (Sigma-Aldrich C8106-500G). Under a dissecting microscope, the vertebral bodies were resected to expose the underlying spinal cord from cervical through sacral segments, and the spinal cord extracted with dorsal and ventral roots. The dura mater was removed from the cord and the tissue transferred to an incubation chamber filled with warmed (28-30°C), continuously bubbled artificial cerebrospinal fluid (aCSF) that contained (mMol/L in 18.2 MΩ H_2_O): sodium chloride (111; Sigma-Aldrich S7653), potassium chloride (3.085; Sigma-Aldrich P5405), d-glucose (10.99; Sigma-Aldrich 158968), sodium bicarbonate (25; Sigma-Aldrich S5761), magnesium sulfate heptahydrate (1.26; Sigma-Aldrich 63145), calcium chloride (2.52; Sigma-Aldrich C8106), and potassium phosphate monobasic (1.1; Sigma-Aldrich P0662). Isolated spinal cords were kept in warmed aCSF for approximately 30 minutes and then switched to room temperature.

#### Recordings

After at least 60 minutes recovery, isolated spinal cords were transferred to a recording chamber and pinned ventral side up. The chamber was continuously perfused with room temperature (20-22°C) carbogen-bubbled aCSF (7.5 rpm flow rate). Pulled and sized single barrel borosilicate glass suction electrodes (WPI 1B120-4) were attached to custom-made bipolar electrode holders and then attached to rostral (bilateral L1 or L2) and caudal (bilateral L4 or L5) lumbar ventral roots and connected to either custom-made or commercially sourced (SuperTech) headstages which were then connected to custom-made amplifiers. Signals were differentially amplified (10,000x), bandpass-filtered (30 Hz to 10kHz), sampled at 10 kHz using Spike2 (CED, v9.04; UK), digitized using an A/D converter (CED Power3a and 1401 Micro3 + ADC12 expansion), and saved for offline analysis.

The recording pipeline consisted of alternating rounds of normal aCSF and drug-induced fictive locomotion. Each preparation underwent at least 3 bouts of normal aCSF (each 10-15 min) and at least 2-3 bouts of drug-induced fictive locomotion (each 30-45 min). In P1-P2 spinal cords, fictive locomotion was elicited using a cocktail of 5 µM N-Methyl-D-aspartic acid (NMDA; Sigma-Aldrich M3262-100mg) and 10 µM serotonin (5-HT, Sigma-Aldrich H9523-100mg)^74^. For P3-P5 spinal cords, the cocktail was supplemented with 40 µM dopamine (Sigma-Aldrich H8502-5g)^74^. Files were annotated with timestamps to denote when drugs were applied and subsequently washed out. Spinal cords that showed >30 minutes of stable bursting in at least 3 ventral roots were selected for prescreening and subsequent analyses.

#### Analysis

Starting at the washout timestamp, a retro-sliding 180s window was used to segment the data into 3-minute epochs until bursting onset was observed. After segmentation, the first and last epochs from all bouts were excluded. Segmented data were then processed to remove, where needed, signal glitches using the Spike2 “FixGlitch” script. Signals were then down-sampled to 5 kHz, imported into SpinalCoreN MATLAB-based software^38^, full-wave rectified, and high-pass filtered. Bursting activity from all ventral roots, bouts, and intra-bout 3-minute epochs were processed using a set of wavelet transformations. First, we plotted the power-frequency spectrum of the signal over time for each 3-minute bout. Next, we set a minimum power threshold of 128 to isolate signal from background noise. Empirical testing of multiple datasets confirms this cut-off eliminates the ultralow power, high frequency signal contaminants embedded within the background. We then set a blanket region of interest (ROI) from x_1_=0 sec to x_2_=180 sec, y_1_=0 Hz to y_2_=4 Hz to capture the total power-frequency spectrum. Within this blanket ROI, we then isolated the signal within the dominant power-frequency band(s) using a minimum power threshold of 512 (4x the minimum signal:noise cut-off). After high-power ROI detection, the signal burst frequency and cycle duration over time were extracted for each ventral root and subsequently analysed. Once the dominant power-frequency bands were detected in each ventral root, we then performed a set of cross-wavelet transformations focusing on the following lumbar ventral root pairs: bilateral rostral (L1-L2), bilateral caudal (L4-L5), and ipsilateral (L1/L2-L4/L5). Using the same tiered ROI strategy, we isolated the shared dominant power-frequency bands and extracted the cross-root burst frequency, cycle duration, phase relationship, and coherence over time for each epoch. Data shown are from N=5 littermate controls (n=8-12 epochs per animal) aged P1-P4 and N=11 spinal *Tor1a* d-cko mice (n=4-13 epochs per animal) aged P2-P5 using the animal as the experimental unit for analysis. N=2 P1 spinal *Tor1a* d-cko mice were excluded from group comparisons as mean outcomes were broadly similar to littermate controls and our focus was on quantifying the circuit disruption(s) associated with the movement disorder (data excluded are shown in Fig. S3).

### *In vitro* electrophysiology: monosynaptic reflex

#### Preparation

*In vitro* monosynaptic reflex testing was performed following a modified protocol previously described^75^. N=24 littermate control and N=21 spinal *Tor1a* d-cko pups aged P7-P13 were anaesthetized via i.p. injection of ketamine:xylazine (300 mg/kg:30 mg/kg; 0.01 ml/g body weight), decapitated, and eviscerated as described above. Then, from the thoracic to sacral level, the spinal cord was rapidly dissected with dorsal and ventral roots attached. Once extracted, the cord was pinned to the dissection chamber ventral side up, longitudinally hemisected through the anterior midline fissure using a fine insect pin (Watkins & Doncaster, E6901 D1 pins), and transferred to an incubation chamber filled with warmed (28-30°C) and continuously bubbled aCSF. Hemisected spinal cords were kept in warmed aCSF for approximately 30 minutes and then switched to room temperature.

#### Recordings

All recordings were performed at room temperature (20-22°C) at least 60 minutes post-dissection. Hemisected spinal cords were transferred to a recording chamber continuously perfused with carbogen-bubbled aCSF and pinned such that the L1-L5 dorsal and ventral roots were accessible. For afferent stimulation, suction electrodes connected to a stimulus isolator (AMPI Isoflex) were attached to the L1-L5 dorsal roots. For recording the evoked monosynaptic reflexes, L1-L5 ventral roots were attached to extracellular recording electrodes connected to custom-made bipolar electrode holders and headstage. Motor response threshold (MT) was determined at the start of each experiment on a root-by-root basis for each individual animal. Dorsal roots were stimulated with a single 0.1ms square pulse of 0.1-0.3 V. MT was defined as the minimum stimulus intensity required to evoke a response above baseline threshold in N=5+/10 sweeps using a 10s interstimulus interval. Threshold was subsequently validated by decreasing the stimulus intensity just below the putative threshold to confirm lack of an evoked response and then increasing the stimulus intensity slightly above the putative threshold to confirm increased amplitude in the monosynaptic reflex. After defining MT, N=10 sweeps/trial/root at inter-stimulus intervals of 30s were recorded at the following multiples of threshold: 1.0x (MT), 1.2x, 1.5x, 2.0x, 5.0x, and 10.0x. In select cases, 10.5x stimulation was used to confirm supramaximal threshold was reached. Signals were differentially amplified (10-10,000x), bandpass-filtered (30 Hz to 10kHz), sampled at 10 kHz using Signal (CED, v6.06), digitized using an A/D converter (CED Power3a and 1401 Micro3 + ADC12 expansion), and saved for offline analysis.

#### Analysis

Monosynaptic reflex data were analysed using a custom-written code for Signal (code is available at https://github.com/Brownstone-lab and our open access UCL Research Data Repository). Evoked response measurements were calculated on a sweep-by-sweep basis relative to each sweep’s baseline, or threshold level of activity. Spike onset was calculated as the time at which mean amplitude (V) exceeded 2S.D. of a 10ms baseline window that preceded the stimulus onset by 5ms. Using this threshold value, active cursors were used to identify the time to rising and falling threshold crossings and peak amplitude of the first excursion in the triphasic reflex waveform. With the TrendPlot function, the following data were extracted from each sweep: (1) latency to onset (time to first rising threshold crossing sustained for more than 0.5ms), (2) latency to peak (time at which the maximally evoked reflex response occurs), (3) offset (time to first falling threshold crossing sustained for more than 0.1ms), (4) response duration (calculated difference: offset - latency to onset), and (5) peak amplitude. Two duration outliers (>3SD) were excluded from analysis for the L5 response duration dataset at P9-P10 (values of 11.4ms and 14.3ms, respectively). Outliers are shown in Fig. S4H.

### *In vitro* electrophysiology: patch-clamp recordings in lumbar motoneurons and conduction velocity estimates

#### Preparation

N=11 littermate control and N=16 spinal *Tor1a* d-cko mice aged P1-P13 were anesthetized with a mixture of ketamine:xylazine (100 mg/kg:10 mg/kg) and decapitated. The vertebral column was removed and pinned ventral side up in a chamber filled with recording aCSF containing (in mM): 113 NaCl, 3 KCl, 25 NaHCO_3_, 1 NaH_2_PO_4_, 2 CaCl_2_, 2 MgCl_2_, and 11 D-glucose continuously bubbled with carbogen. Vertebral bodies were resected and the spinal cord isolated from lower-thoracic to upper sacral segments. For whole cord recordings (P1 only), the cord was transected above the lumbosacral region, with lumbar dorsal or ventral roots slightly trimmed and kept intact. For single-cell recordings, the isolated lumbosacral cord was glued longitudinally to an agar block with the ventral or dorsal side up with intact roots, and an oblique cut above the L3 region was performed to allow visualization of the central canal^76^. The glued cord was then immersed in a vibratome chamber (Leica VT1200) with ice-cold aCSF (~2°C) comprising (in mM): 130 K-gluconate, 15 KCl, 0.05 EGTA, 20 HEPES, 25 D-glucose, 3 Na-kynurenic acid, 2 Na-pyruvate, 3 Myo-inositol, 1 Na-L-ascorbate, pH 7.4 with NaOH^77^. The edge of the vibratome blade was aligned with the central canal, and the tissue was longitudinally sectioned, thus obtaining a dorsal or ventral horn ablated spinal cord preparation with L3-L5 segments preserved. The tissue was transferred to a chamber with recording aCSF for incubation at 37°C for 30-45 minutes, and then kept at room temperature constantly bubbled with carbogen. All experiments were performed in either the dorsal or ventral-horn ablated preparation, with the exception of two experiments performed on P1 mice (1 littermate control and 1 spinal *Tor1a* d-cko), where recordings were performed in the intact spinal cord and motoneurons were targeted in the most ventral nuclei.

#### Recordings

Whole-cell patch clamp recordings were performed using either an Axopatch 200B amplifier or a MultiClamp 700B (Molecular Devices). Signals were filtered at 5 kHz and acquired at 50 kHz using a Digidata 1440A A/D board (Molecular Devices) and Clampex 10 software (Molecular Devices). Glass pipettes of borosilicate glass (GC150F, Harvard Apparatus) were pulled using a P1000 Flaming-Brown puller (Sutter Instruments) and fire polished to a resistance of ~2-4 MΩ. Patch pipettes were filled with an intracellular solution containing (in mM): 125 K-gluconate, 6 KCl, 10 HEPES, 0.1 EGTA, 2 Mg-ATP, pH 7.3 with KOH, and osmolarity of 290-310 mOsm.

Motoneurons distributed near the surface of the ventral and dorsal horn ablated preparations were targeted and visualized using an Eclipse E600FN Nikon microscope (Nikon) equipped with infrared differential interference contrast connected to a digital camera (Nikon DS-Qi1Mc). Motoneuron capacitance and resistance were determined either from the voltage response to a brief 200ms current step (10 to 50pA) in current clamp, or the current change to a 5mV voltage step in voltage clamp. In the ventral horn ablated preparation, dorsal roots were stimulated with an isolated current stimulator (Digitimer DS3) at an intensity of 1.5-3x and 3-5x the threshold for evoking an initial response in the recorded cell. Excitatory postsynaptic currents (EPSCs) were recorded following dorsal root stimulation. In some patch experiments, the conduction velocity observed from the whole root stimulation was compared with that measured from stimulation of a subset of fibres. This was achieved by stretching the stimulated root and placing a second extracellular stimulation electrode in different positions along the root. The size of the second stimulation electrode was just sufficient to “pinch” the root (typically 20-40μm) and therefore we could evoke spikes only in the subset of fibers close to the electrode. Microstimulations were delivered at 1x threshold, in order to minimize the number of activated fibers. For extracellular recordings of afferent and efferent volleys, root responses were acquired with an NPI Ext-02F amplifier and the stimulating and recording electrodes were placed along a stretched dorsal or ventral root, allowing a more reliable measurement of the conduction velocity.

#### Analysis

The conductance of the dorsal root-evoked EPSCs was calculated from the size of the current recorded at a holding voltage of −60mV and assuming a reversal of 0mV for excitatory conductances. In order to assess the contribution of the synaptic conductance to the overall resting conductance of each cells, values of the synaptic conductance are reported scaled to the conductance of each cell. The latency was measured from the onset of the stimulus artefact to the point at which the trace crossed a threshold of 2x the S.D. of the baseline noise. The charge transfer following a dorsal root stimulation was measured by taking the integral of the response in a window ranging from the end of the stimulus artefact and extending for 50-200ms. The latency of the response from the motoneuron and estimated (using ImageJ) distance from microstimulating point to the recorded motoneuron were used to get an estimation of the conduction velocity in patch recordings. This procedure systematically overestimates the conduction velocity, since the measured distance between recorded cell and stimulation point does not account for the length of the fibre in the dorso-ventral direction from the entry point of the root to the recorded cell. Raw traces were analysed with Clampfit (v10) software.

### Statistical analysis

Statistical analyses were performed using Origin Lab (v2021), Microsoft Excel, Data Analysis with Bootstrap-coupled ESTimation Python script (DABEST)^78^, MATLAB (R2021a), MATLAB-based SpinalCoreN (v2018b), GraphPad Prism (v9), or SigmaPlot (v22). Differences between groups were considered statistically significant at p≤0.05. Experimental units are defined in figure legends. Two-tailed p-values are reported. Outliers, which were defined as any value that exceeded average ± 3SD, were excluded from datasets prior to analysis.

#### Normality and equality of variance

Data were tested for normal distribution using the Shaprio-Wilk test. Equality of variance was tested using Brown-Forysthe ANOVA (for non-normally distributed data), Levene’s test (for normally distributed data), Welch’s test (reporting F-statistic), or F-test.

#### Mean comparisons

Means were compared using either paired t-tests, independent t-tests with or without Welch’s correction for unequal variance, Mann-Whitney non-parametric test, or two-way ANOVAs followed by Tukey’s *post hoc* t-test with p-value correction for multiple comparisons. Regression and slope analyses were performed for the lines of best fit for age-dependent changes in DR-evoked EPSC area and latency using Fisher’s r to z transformation.

#### Circular statistics

Burst coordination datasets were analysed using Watson’s non-parametric two-sample U^2^ test as previously described^79,80^.

#### Data presentation and estimation statistics

Group data are shown as box plots that include the minimum, first quartile, median, third quartile, and maximum values. Individual mean values are shown as filled circles overlaid onto box plots, where relevant. Raw data are plotted as small dots adjacent to box plots. Mean differences are shown using Gardner-Altman estimation plots showing the effect size from a sampling distribution, 95% confidence interval (CI) and mean obtained with bootstrapping (resampling done 5,000-10,000x)^78^.

## Supplemental information

**Figure S1.**
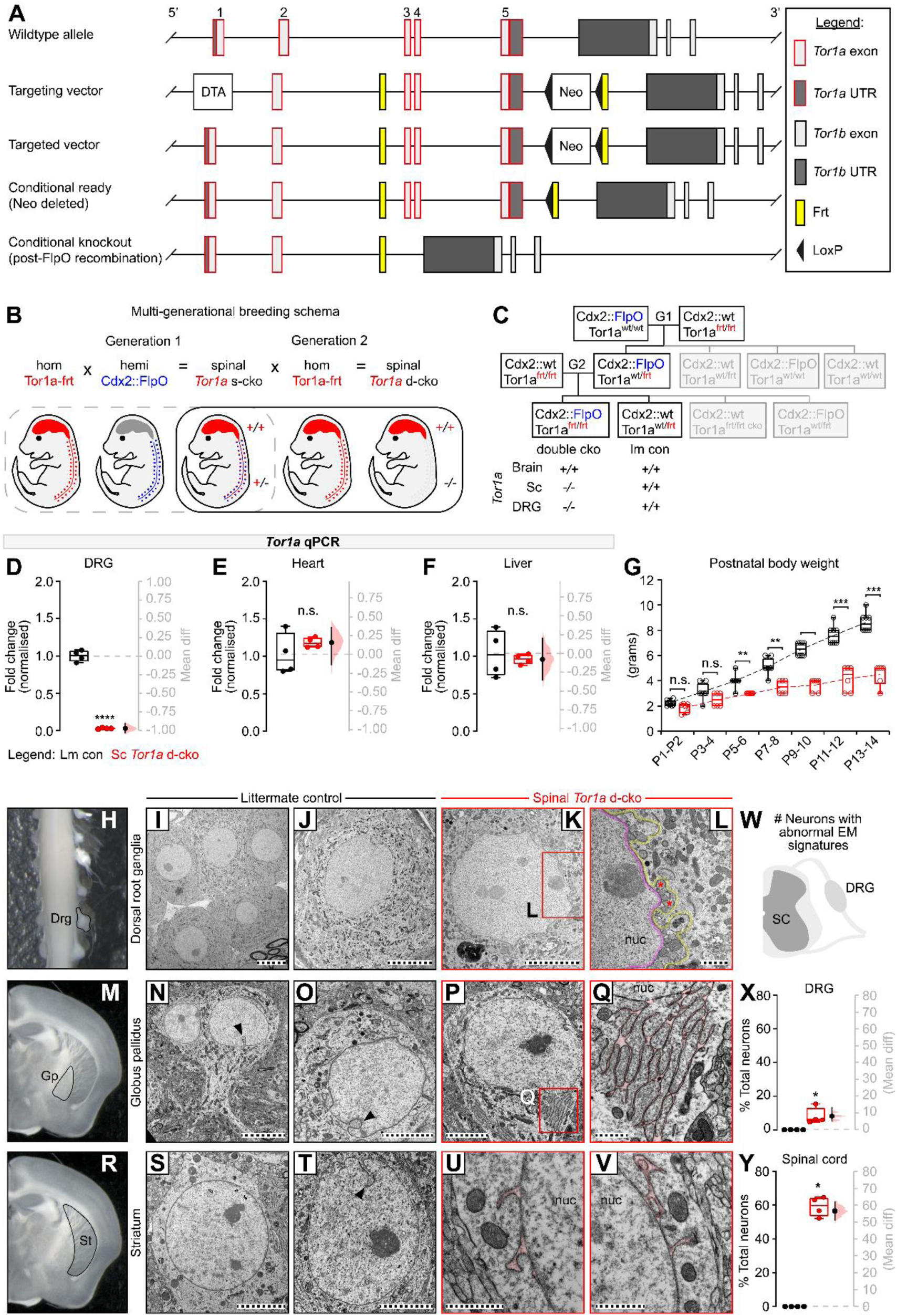
Tor1a-frt model details and breeding strategy to generate spinal-restricted *Tor1a* d-cko mice. (**A**) Targeting strategy to generate the Tor1a-frt mouse. (**B-C**) Multi-generational breeding schema to produce a biallelic “double” conditional knockout (d-cko) of *Tor1a* in the spinal cord and DRGs. (**D-F**) qPCR confirming the site-specificity of the *Tor1a* conditional knockout. (**G**) Postnatal body weights are reduced in spinal *Tor1a* d-cko mice as compared to littermate controls, but mice still gain weight over time. (**H-Y**) Ultrastructural findings. In littermate controls, neurons in the lumbar spinal dorsal root ganglia (**H-J**), globus pallidus (**M-O**), and striatal nuclei (**R-T**) show normal nuclear envelope (NE) morphology, including occasional nucleoplasmic reticulations (arrowheads). (**K-L**) In contrast to the homonymous lumbar segment (Fig. 1), few DRG neurons show nuclear envelope vesiculation and other ultrastructural signatures of torsinA dysfunction. In all neurons screened, no nuclear envelope-derived vesicles were detected in globus pallidus (**P-Q**) or striatal (**U-V**) neurons of spinal *Tor1a* d-cko mice. However, instances of outer nuclear membrane protrusions and separation between the nuclear bilayers were noted (highlighted), sometimes forming complex branches (**P-Q**). (**X**) Percent total DRG neurons and (**Y**) lumbar spinal cord neurons that show morphological abnormalities in P18 littermate control (N=4) and spinal *Tor1a* d-cko mice (N=4). \**P*<0.05, Mann-Whitney test. Group data shown (box plots) with individual values overlaid (circles) and mean differences (estimation plots). Statistics details in Table S1. Scale bars: 10µm (I-J, N), 5µm (K, O-P, S-T), 1µm (L, Q, U-V). Nuc = nucleus. All ultrastructure images generated in this study available at our open access UCL Research Data Repository. (Box plots) Data shown are group average with individual values overlaid (circles). Mean differences shown as Gardner-Altman estimation plots. Statistics details in Table S1. Associated with Figure 1.

**Figure S2.**
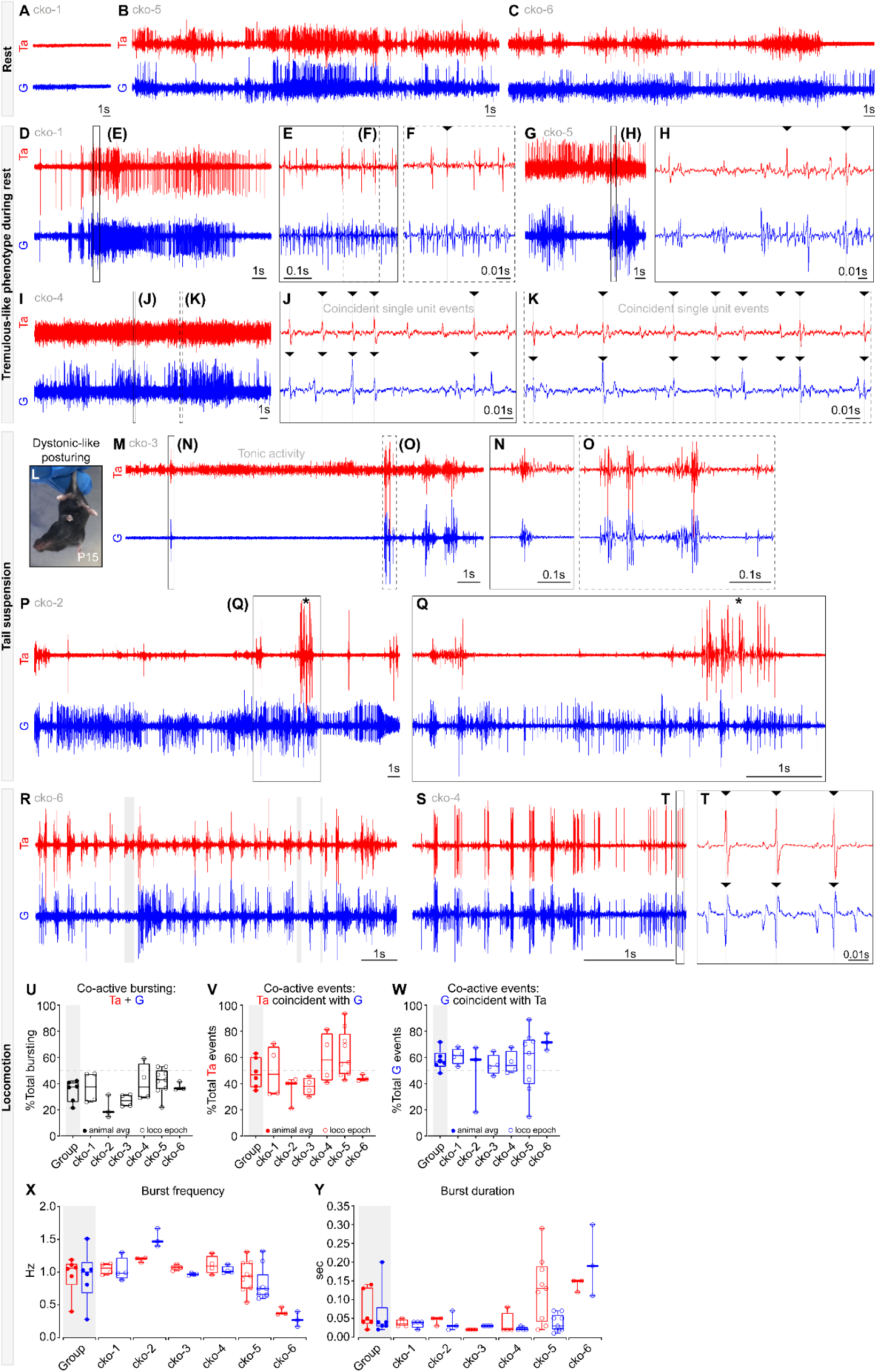
Abnormal hindlimb EMG activity in spinal *Tor1a* d-cko mice. (**A-C**) Representative examples of tibialis anterior (Ta) and gastrocnemius (G) activity at rest. (**D-K**) During rest, there was the incursion of a tremulous-like phenotype marked by increased EMG activity and coincident single unit spikes. (**L-Q**) EMG activity observed during tail suspension included tonic activity, high amplitude bursting, and co-contractions. (**R-S**) EMG bursting observed during locomotion. (**U**) Group and per animal co-contractions observed of total bursting activity. (**V-W**) Total EMG co-activity observed, inclusive of high frequency single unit bursts. Data shown include: (**V**) the proportion of Ta events that are co-active with G, and (**W**) proportion of G events that are co-active with Ta. (**X**) Group and individual animal burst frequency and (**Y**) burst duration observed during locomotion. “Group”: group mean ± SD with individual values overlaid (circles). “cko-1 to -6”: individual mean ± SD for total bouts analysed/animal with individual bout values overlaid (circles). Asterisks: bursts were truncated for visual representation. Associated with Figure 2.

**Figure S3.**
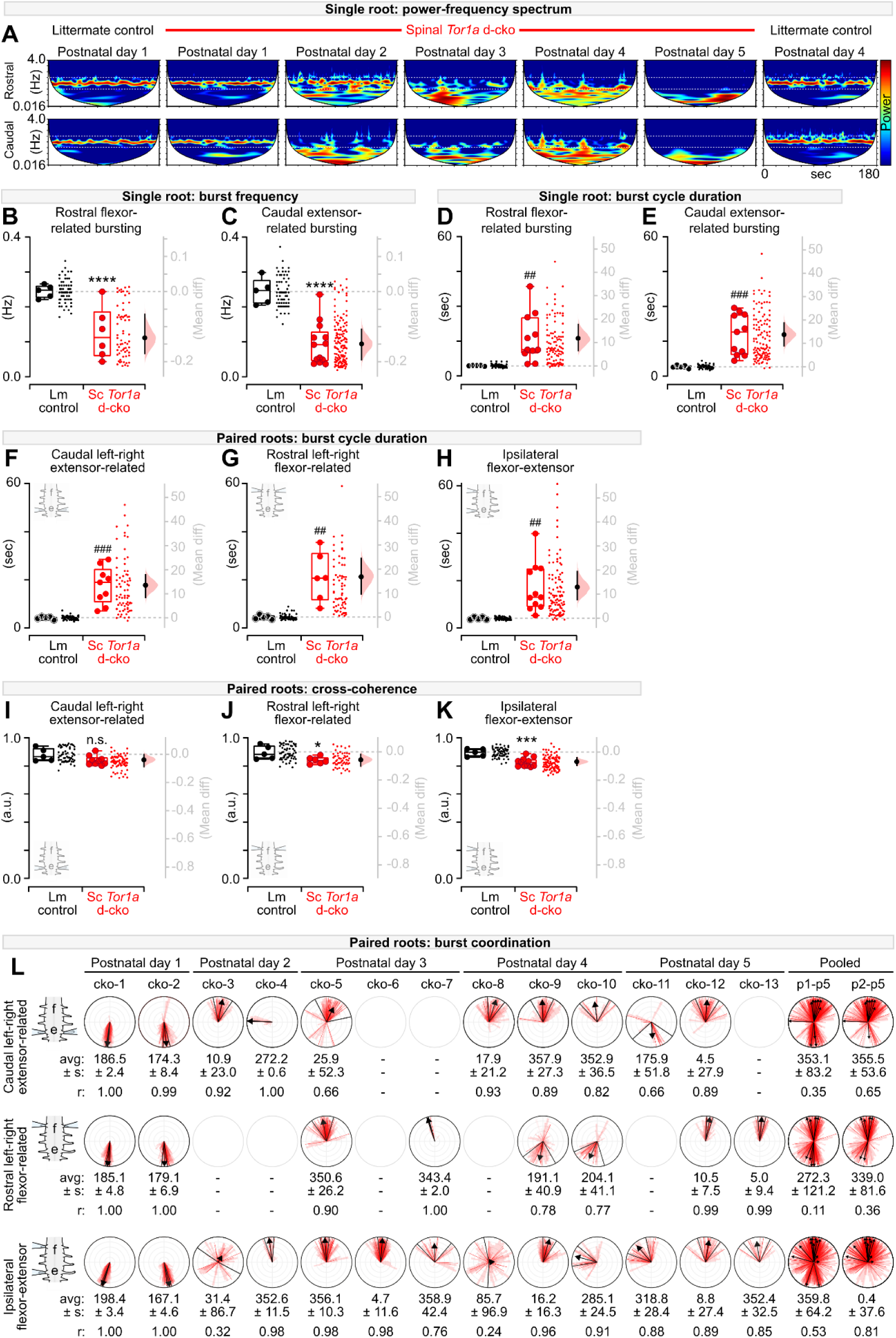
Spinal-restricted *Tor1a* deletion leads to disorganised motor output. (**A**) Single root wavelet transformations. Power range (cool-warm colours): 2^7^-2^13^. (**B-E**) Single root burst analysis, control (*N*=5) vs spinal *Tor1a* d-cko mice(*N*=6-11). (**B**) Rostral burst frequency: 0.24±0.02Hz vs 0.08±0.04Hz; \*\*\*\**P*<0.0001, independent t-test. (**C**) Caudal: 0.24±0.04Hz vs 0.08±0.04Hz; \*\*\*\**P*<0.0001. (**D**) Rostral cycle duration: 4.65±0.17s vs 16.44±10.28s; ^##^*P*<0.01, independent t-test with Welch’s correction. (**E**) Caudal: 4.26±0.64s vs 18.01±8.70s; ^###^*P*<0.001. (**F-K**) Cross-root burst analysis, control vs spinal *Tor1a* d-cko mice. (**F**) Caudal left-right cycle duration: 4.20±0.57s vs 17.77±7.65s; ^###^*P*<0.001. (**G**) Rostral: 4.51±0.84s vs 21.25±10.21s; ^##^*P*=0.01. (**H**) Ipsilateral rostrocaudal: 4.06±0.54s vs 17.00±10.35s; ^##^*P*<0.01. (**I**) Caudal cross-coherence: 0.88±0.05 vs 0.84±0.03; *P*=0.09, independent t-test. (**J**) Rostral: 0.90±0.05 vs 0.84±0.02; \**P*<0.05. (**K**) Ipsilateral rostrocaudal: 0.89±0.03 vs 0.83±0.03; \*\*\**P*<0.001. (**L**) Burst coordination in spinal *Tor1a* d-cko mice. Mean phase ± circular variance (s) and r-value reported. Clockwise from top of circular plot=0° (synchrony), bottom=180° (alternation). (Box plots) Data shown are group average with individual values overlaid (circles). Adjacent dots: epochs analysed/animal. Mean differences shown as estimation plots. (Circular plots) Bold arrows: orientation=mean phase, length (0-1, concentric gray circles)=concentration of observations, black wedge=circular variance. Group data overlaid onto total observations from all epochs (wedges) and epoch-specific averages (lines). Statistics details in Table S1. Associated with Figure 3.

**Figure S4.**
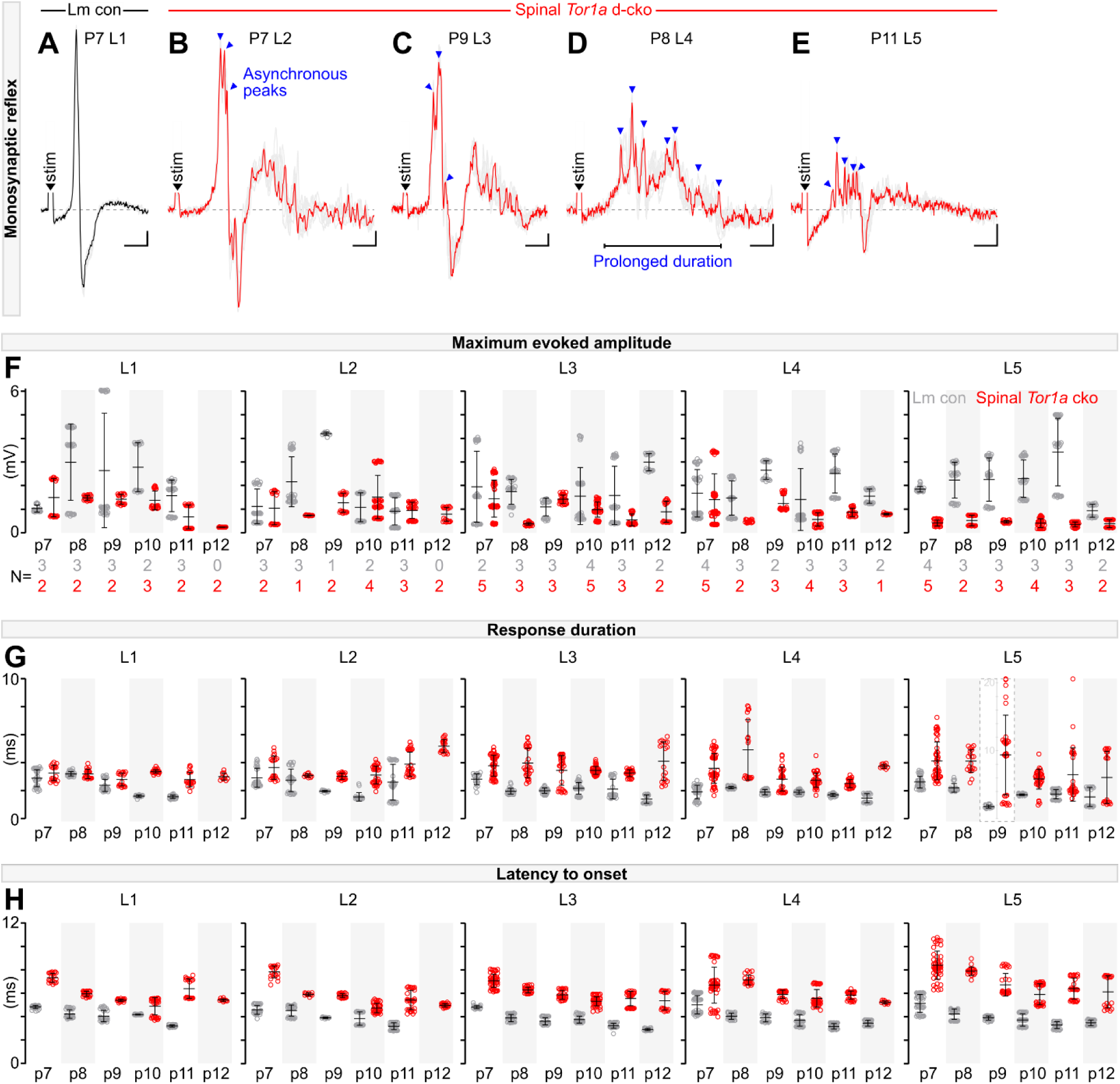
*Tor1a*-deleted spinal circuits have abnormal monosynaptic reflexes. (**A**) Representative example of monosynaptic reflex in littermate control. (**B-E**) Monosynaptic reflexes in spinal *Tor1a* d-cko mice across space (L2-L5) and age (P7-P11) show increased latency to onset, prolonged duration, and the infiltration of multiple asynchronous peaks. *N*=10 sweeps shown/root at 2x motor threshold. One sweep is highlighted to illustrate triphasic waveform observed. (**F-H**) Per animal per segment per timepoint data shown for pooled outcomes reported in Fig. 4. (**F**) Maximum evoked amplitude in monosynaptic reflexes observed in littermate controls (grey) and spinal *Tor1a* d-cko mice (red). Data shown include mean ± S.D. (lines) overlaid onto *N*=10 reflex responses analysed per animal (circles). Number of animals analysed per root per time point reported below graphs. (**G**) Monosynaptic response duration observed at 2x motor threshold. Note scale change for P9 L5 data to accommodate 2 outliers. (**H**) Latency to onset was increased in spinal *Tor1a* d-cko mice compared to controls. Scale: x=5ms, y= 0.1mV (A, B) or 0.05mV (C-E). Associated with Figure 4.

**Figure S5.**
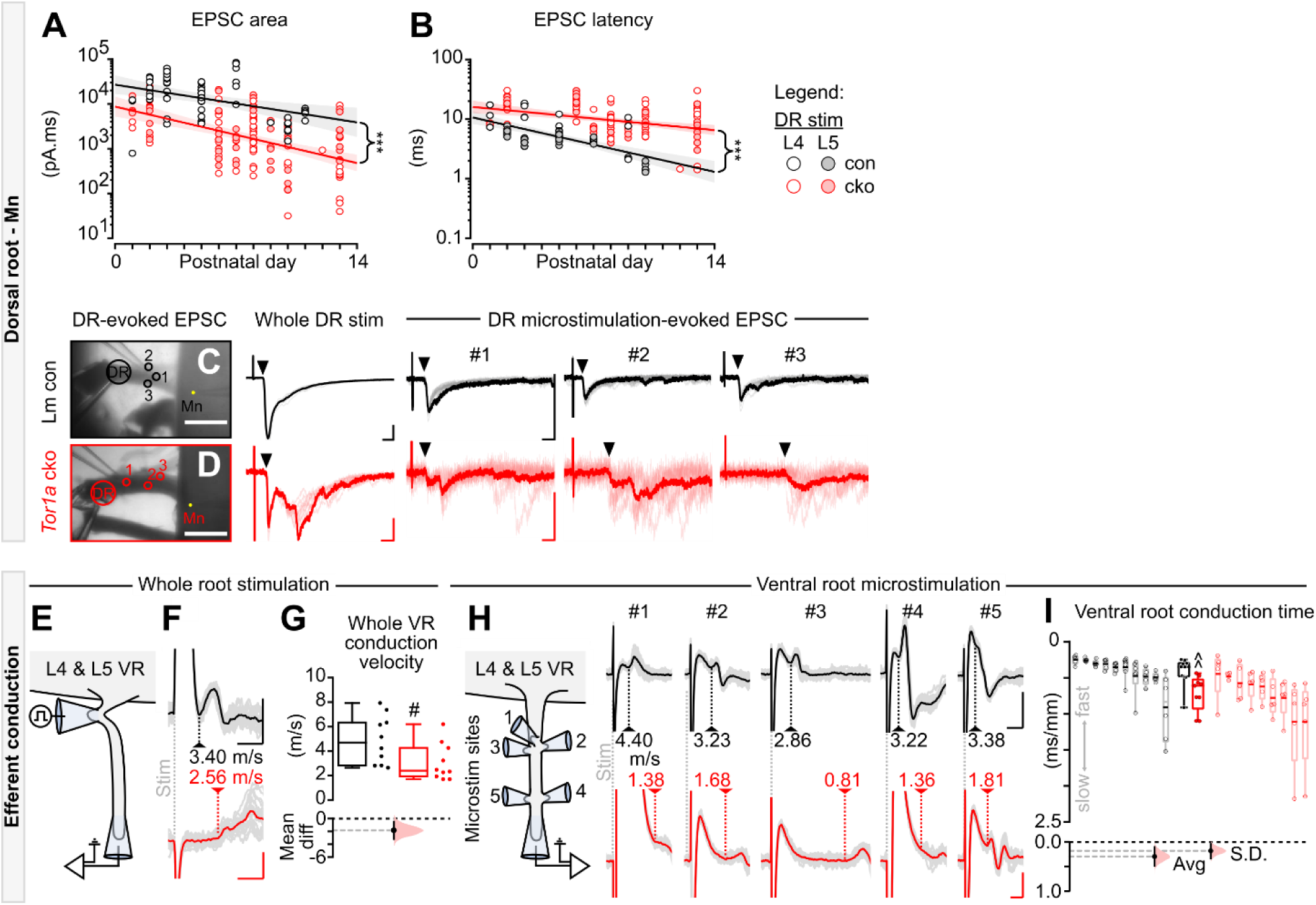
Spinal *Tor1a* d-cko leads to changes in all components of the monosynaptic reflex. Log plots illustrating regression analysis and slope comparisons of group-related, age-dependent changes in (**A**) DR-evoked EPSC area (P1-P13 *N*=61 control vs *N*=130 spinal *Tor1a* d-cko mice; Y=-1.6e3x+2.9e4 with R^2^=0.066 vs Y=-6.9e2x+9.3e3 with R^2^=0.202, \*\*\**P*<0.001, Fisher’s r to Z test) and (**B**) EPSC latency (*N*=60 control vs *N*=131 spinal *Tor1a* d-cko mice; Y=-0.7x+9.6 with R^2^=0.242 vs Y=-0.6x+16.5 with R^2^=0.108, \*\*\**P*<0.001). Circles: individual L4 (unfilled) or L5 (filled) DR-evoked responses in control (gray) or *Tor1a* d-cko (red) mice. Lines of best fit overlaid onto 95% confidence intervals. (**C-D**) DR microstimulation-evoked EPSC reveals increased latency and asynchronous activation of motoneurons in spinal *Tor1a* d-cko mice (P7 control, P9 spinal *Tor1a* d-cko mice). Scales: DIC=500µm; black scales (whole, micro): x=5ms, y=500pA; red: x=5ms, y=100pA (whole), y=50pA (micro). (**E**) L5 ventral root (VR) microstimulation and recording of efferent volleys. (**F**) Representative examples of microstimulation-evoked efferent volleys at threshold (P8 control, P7 spinal *Tor1a* d-cko mice). Vertical line: first deflection point used to estimate conduction velocity (indicated). Scale: x=0.5ms, y=0.025mV. (**G**) L4 & L5 VR conduction velocity is reduced in spinal *Tor1a* d-cko (P6-P9 *N*=10 control and *N*=10 spinal *Tor1a* d-cko ventral roots, ^#^*P*<0.05, independent t-test). (**H**) Representative examples of microstimulation-evoked efferent volleys (P8 control, P7 spinal *Tor1a* d-cko mice). Vertical line: first deflection point used to estimate conduction velocity (indicated). Black scale: x=0.5ms, y=0.05mV vs red x=1ms, y=0.025mV. (**I**) Efferent volleys in spinal *Tor1a* d-cko mice showed longer conduction times (P6-P9 N=10 control vs N=9 cko, average, ^^P<0.01, Mann-Whitney test) and increased variances in conduction times (P<0.05). Averaged data shown in black and red box plots with individual values (means within each root) overlaid (filled circles) and mean differences (estimation plots). The responses from individual roots - including raw (open circles) and averaged (bold line) values - are shown as de-saturated box plots adjacent to the averaged dataset. Statistics details in Table S1. Associated with Figure 5.

**Movie S1.** Representative postnatal video recordings of littermate control (#1161022) vs spinal *Tor1a* d-cko mice (#1161023) showing onset and progression of dystonic-like signs over postnatal maturation. All videos assessed are available at our open access UCL Research Data Repository.

**Movie S2.** Tail suspension-induced paw clasping and clenching in spinal *Tor1a* d-cko mice.

**Movie S3.** Range of discoordinated limb movements observed in pre-weaned spinal *Tor1a* d-cko mice.

**Movie S4.** Range of trunk torsion and instability observed throughout postnatal maturation in spinal *Tor1a* d-cko mice.

**Movie S5.** Range of abnormal postures observed at rest and during voluntary movements in spinal *Tor1a* d-cko mice.

**Movie S6.** Intrusion of tremulous-like phenotype in spinal *Tor1a* d-cko mice.

**Movie S7.** Confining the biallelic conditional knockout to dorsal root ganglia (DRG) neurons (Avil^wt/cre^;Tor1a^flox/flox^) does not produce early onset generalised torsional dystonia.

## Funding

Wellcome Trust Investigator Grant 110193 (RMB)

Medical Research Council Research Grant MR/V003607/1 (RMB) and MR/R011494 (MB and RMB)

Biotechnology and Biological Sciences Research Council Research Grant BB/S005943/1 (MB)

European Molecular Biology Organisation Long Term Fellowship ALTF 495-2018 (AMP)

Sir Henry Wellcome Postdoctoral Fellowship 221610/Z/20/Z (FN)

Royal Society Newton International Fellowship NIF\R1\192316 (MGO)

Human Frontiers Science Program Long Term Fellowship LT000220/2017-L (SS)

Medical Research Council Core Funding Grant MC/U12266B (IW)

Wellcome Trust Facility Grant 218278/Z/19/Z (IW)

## Resource availability

Data and custom-written code are available at our UCL Research Data Repository entitled “Pocratsky et al 2022” (see methods for access details). The Tor1a-frt mouse model generated in this study is available through the MMRRC commercial mouse repository. For the purpose of Open Access, the authors will apply a CC by public copyright licence to the accepted manuscript. Further information and requests for resources and reagents should be directed to the lead contact, Rob Brownstone.

## Notes

### Summary of Updates

Additional supplemental data were provided regarding the conditional knockout of Tor1a in dorsal root ganglia neurons. Additional control data was generated for EMG recordings.

## References

1. Saxena, S., Russo, A.A., Cunningham, J., and Churchland, M.M. (2022). Motor cortex activity across movement speeds is predicted by network-level strategies for generating muscle activity. Elife 11. 10.7554/eLife.67620.

2. Lee, J., Wang, W., and Sabatini, B.L. (2020). Anatomically segregated basal ganglia pathways allow parallel behavioral modulation. Nat Neurosci 23, 1388–1398. 10.1038/s41593-020-00712-5.

3. Klaus, A., Silva, J.A.d., and Costa, R.M. (2019). What, If, and When to Move: Basal Ganglia Circuits and Self-Paced Action Initiation. 42, 459–483. 10.1146/annurev-neuro-072116-031033.

4. MacLean, J.N., Watson, B.O., Aaron, G.B., and Yuste, R. (2005). Internal dynamics determine the cortical response to thalamic stimulation. Neuron 48, 811–823. 10.1016/j.neuron.2005.09.035.

5. Thanawalla, A.R., Chen, A.I., and Azim, E. (2020). The Cerebellar Nuclei and Dexterous Limb Movements. Neuroscience 450, 168–183. 10.1016/j.neuroscience.2020.06.046.

6. Brownstone, R.M., and Chopek, J.W. (2018). Reticulospinal Systems for Tuning Motor Commands. Front Neural Circuits 12, 30. 10.3389/fncir.2018.00030.

7. Kiehn, O. (2006). Locomotor circuits in the mammalian spinal cord. Annu Rev Neurosci 29, 279–306. 10.1146/annurev.neuro.29.051605.112910.

8. Grütz, K., and Klein, C. (2021). Dystonia updates: definition, nomenclature, clinical classification, and etiology. Journal of Neural Transmission 128, 395–404. 10.1007/s00702-021-02314-2.

9. Rothwell, J.C., Obeso, J.A., Day, B.L., and Marsden, C.D. (1983). Pathophysiology of dystonias. Adv Neurol 39, 851–863.

10. Cohen, L.G., and Hallett, M. (1988). Hand cramps: clinical features and electromyographic patterns in a focal dystonia. Neurology 38, 1005–1012.

11. Novikova, V.P. (1981). Spinal mechanisms of motor disturbances in torsion dystonia (an electromyographic analysis). Neuroscience and behavioral physiology 11, 353–357. 10.1007/bf01184199.

12. Koelman, J.H.T.M., Willemse, R.B., Bour, L.J., Hilgevoord, A.A.J., Speelman, J.D., and Visse, B.W.O.d. (1995). Soleus H-reflex tests in dystonia. Movement Disorders 10, 44–50. 10.1002/mds.870100109.

13. Meyer, A., and Hierons, R. (1965). ON THOMAS WILLIS’S CONCEPTS OF NEUROPHYSIOLOGY. I. Medical history 9, 1–15. 10.1017/s002572730003009x.

14. Jinnah, H.A., and Hess, E.J. (2017). Evolving concepts in the pathogenesis of dystonia. Parkinsonism & related disorders. 10.1016/j.parkreldis.2017.08.001.

15. Neychev, V.K., Gross, R.E., Lehericy, S., Hess, E.J., and Jinnah, H.A. (2011). The functional neuroanatomy of dystonia. Neurobiol Dis 42, 185–201. 10.1016/j.nbd.2011.01.026.

16. Newby, R.E., Thorpe, D.E., Kempster, P.A., and Alty, J.E. (2017). A History of Dystonia: Ancient to Modern. Movement disorders clinical practice 4, 478–485. 10.1002/mdc3.12493.

17. Paudel, R., Hardy, J., Revesz, T., Holton, J.L., and Houlden, H. (2012). Review: genetics and neuropathology of primary pure dystonia. Neuropathology and applied neurobiology 38, 520–534. 10.1111/j.1365-2990.2012.01298.x.

18. Sharma, N., Baxter, M.G., Petravicz, J., Bragg, D.C., Schienda, A., Standaert, D.G., and Breakefield, X.O. (2005). Impaired motor learning in mice expressing torsinA with the DYT1 dystonia mutation. J Neurosci 25, 5351–5355. 10.1523/jneurosci.0855-05.2005.

19. Scott, B.L., and Jankovic, J. (1996). Delayed-onset progressive movement disorders after static brain lesions. Neurology 46, 68–74. 10.1212/wnl.46.1.68.

20. Lozano, A.M., Hutchison, W.D., and Kalia, S.K. (2017). What Have We Learned About Movement Disorders from Functional Neurosurgery? 40, 453–477. 10.1146/annurev-neuro-070815-013906.

21. Pappas, S.S., Leventhal, D.K., Albin, R.L., and Dauer, W.T. (2014). Mouse models of neurodevelopmental disease of the basal ganglia and associated circuits. Current topics in developmental biology 109, 97–169. 10.1016/b978-0-12-397920-9.00001-9.

22. Brownstone, R.M. (2020). Key Steps in the Evolution of Mammalian Movement: A Prolegomenal Essay. Neuroscience 450, 135–141. https://doi.org/10.1016/j.neuroscience.2020.05.020.

23. Ozelius, L.J., Page, C.E., Klein, C., Hewett, J.W., Mineta, M., Leung, J., Shalish, C., Bressman, S.B., de Leon, D., Brin, M.F., et al. (1999). The TOR1A (DYT1) gene family and its role in early onset torsion dystonia. Genomics 62, 377–384. 10.1006/geno.1999.6039.

24. Britz, O., Zhang, J., Grossmann, K.S., Dyck, J., Kim, J.C., Dymecki, S., Gosgnach, S., and Goulding, M. (2015). A genetically defined asymmetry underlies the inhibitory control of flexor-extensor locomotor movements. Elife 4. 10.7554/eLife.04718.

25. Pappas, S.S., Darr, K., Holley, S.M., Cepeda, C., Mabrouk, O.S., Wong, J.-M.T., LeWitt, T.M., Paudel, R., Houlden, H., Kennedy, R.T., et al. (2015). Forebrain deletion of the dystonia protein torsinA causes dystonic-like movements and loss of striatal cholinergic neurons. eLife 4, e08352. 10.7554/eLife.08352.

26. Tanabe, L.M., Liang, C.C., and Dauer, W.T. (2016). Neuronal Nuclear Membrane Budding Occurs during a Developmental Window Modulated by Torsin Paralogs. Cell reports 16, 3322–3333. 10.1016/j.celrep.2016.08.044.

27. Pappas, S.S., Li, J., LeWitt, T.M., Kim, J.K., Monani, U.R., and Dauer, W.T. (2018). A cell autonomous torsinA requirement for cholinergic neuron survival and motor control. Elife 7. 10.7554/eLife.36691.

28. Goodchild, R.E., and Dauer, W.T. (2004). Mislocalization to the nuclear envelope: an effect of the dystonia-causing torsinA mutation. Proc Natl Acad Sci U S A 101, 847–852. 10.1073/pnas.0304375101.

29. Goodchild, R.E., Kim, C.E., and Dauer, W.T. (2005). Loss of the dystonia-associated protein torsinA selectively disrupts the neuronal nuclear envelope. Neuron 48, 923–932. 10.1016/j.neuron.2005.11.010.

30. Weisheit, C.E., and Dauer, W.T. (2015). A novel conditional knock-in approach defines molecular and circuit effects of the DYT1 dystonia mutation. Hum Mol Genet 24, 6459–6472. 10.1093/hmg/ddv355.

31. Breakefield, X.O., Kamm, C., and Hanson, P.I. (2001). TorsinA: movement at many levels. Neuron 31, 9–12. 10.1016/s0896-6273(01)00350-6.

32. Bressman, S.B., de Leon, D., Kramer, P.L., Ozelius, L.J., Brin, M.F., Greene, P.E., Fahn, S., Breakefield, X.O., and Risch, N.J. (1994). Dystonia in Ashkenazi Jews: clinical characterization of a founder mutation. Ann Neurol 36, 771–777. 10.1002/ana.410360514.

33. Shashidharan, P., Sandu, D., Potla, U., Armata, I.A., Walker, R.H., McNaught, K.S., Weisz, D., Sreenath, T., Brin, M.F., and Olanow, C.W. (2004). Transgenic mouse model of early-onset DYT1 dystonia. Human Molecular Genetics 14, 125–133. 10.1093/hmg/ddi012.

34. Liang, C.C., Tanabe, L.M., Jou, S., Chi, F., and Dauer, W.T. (2014). TorsinA hypofunction causes abnormal twisting movements and sensorimotor circuit neurodegeneration. J Clin Invest 124, 3080–3092. 10.1172/jci72830.

35. Zhou, X., Wang, L., Hasegawa, H., Amin, P., Han, B.X., Kaneko, S., He, Y., and Wang, F. (2010). Deletion of PIK3C3/Vps34 in sensory neurons causes rapid neurodegeneration by disrupting the endosomal but not the autophagic pathway. Proc Natl Acad Sci U S A 107, 9424–9429. 10.1073/pnas.0914725107.

36. Whelan, P.J. (2003). Developmental Aspects of Spinal Locomotor Function: Insights from Using the in vitro Mouse Spinal Cord Preparation. 553, 695–706. 10.1113/jphysiol.2003.046219.

37. Bonnot, A., Morin, D., and Viala, D. (1998). Genesis of spontaneous rhythmic motor patterns in the lumbosacral spinal cord of neonate mouse. Developmental Brain Research 108, 89–99. https://doi.org/10.1016/S0165-3806(98)00033-9.

38. Mor, Y., and Lev-Tov, A. (2007). Analysis of rhythmic patterns produced by spinal neural networks. J Neurophysiol 98, 2807–2817. 10.1152/jn.00740.2007.

39. Paudel, R., Kiely, A., Li, A., Lashley, T., Bandopadhyay, R., Hardy, J., Jinnah, H.A., Bhatia, K., Houlden, H., and Holton, J.L. (2014). Neuropathological features of genetically confirmed DYT1 dystonia: investigating disease-specific inclusions. Acta Neuropathologica Communications 2, 159. 10.1186/s40478-014-0159-x.

40. Weisheit, C.E., and Dauer, W.T. (2015). A novel conditional knock-in approach defines molecular and circuit effects of the DYT1 dystonia mutation. Human molecular genetics 24, 6459–6472. 10.1093/hmg/ddv355.

41. Paracka, L., Wegner, F., Blahak, C., Abdallat, M., Saryyeva, A., Dressler, D., Karst, M., and Krauss, J.K. (2017). Sensory Alterations in Patients with Isolated Idiopathic Dystonia: An Exploratory Quantitative Sensory Testing Analysis. Frontiers in Neurology 8. 10.3389/fneur.2017.00553.

42. Fiorio, M., Gambarin, M., Valente, E.M., Liberini, P., Loi, M., Cossu, G., Moretto, G., Bhatia, K.P., Defazio, G., Aglioti, S.M., et al. (2006). Defective temporal processing of sensory stimuli in DYT1 mutation carriers: a new endophenotype of dystonia? Brain 130, 134–142. 10.1093/brain/awl283.

43. Albanese, A., Bhatia, K., Bressman, S.B., Delong, M.R., Fahn, S., Fung, V.S., Hallett, M., Jankovic, J., Jinnah, H.A., Klein, C., et al. (2013). Phenomenology and classification of dystonia: a consensus update. Mov Disord 28, 863–873. 10.1002/mds.25475.

44. Tempel, L.W., and Perlmutter, J.S. (1993). Abnormal cortical responses in patients with writer’s cramp. Neurology 43, 2252–2257. 10.1212/wnl.43.11.2252.

45. Kaji, R., Rothwell, J.C., Katayama, M., Ikeda, T., Kubori, T., Kohara, N., Mezaki, T., Shibasaki, H., and Kimura, J. (1995). Tonic vibration reflex and muscle afferent block in writer’s cramp. Ann Neurol 38, 155–162. 10.1002/ana.410380206.

46. Leis, A.A., Dimitrijevic, M.R., Delapasse, J.S., and Sharkey, P.C. (1992). Modification of cervical dystonia by selective sensory stimulation. J Neurol Sci 110, 79–89.

47. Kariminejad, A., Dahl-Halvarsson, M., Ravenscroft, G., Afroozan, F., Keshavarz, E., Goullée, H., Davis, M.R., Faraji Zonooz, M., Najmabadi, H., Laing, N.G., and Tajsharghi, H. (2017). TOR1A variants cause a severe arthrogryposis with developmental delay, strabismus and tremor. Brain : a journal of neurology 140, 2851–2859. 10.1093/brain/awx230.

48. Reichert, S.C., Gonzalez-Alegre, P., and Scharer, G.H. (2017). Biallelic TOR1A variants in an infant with severe arthrogryposis. Neurol Genet 3, e154–e154. 10.1212/NXG.0000000000000154.

49. Isik, E., Aykut, A., Atik, T., Cogulu, O., and Ozkinay, F. (2019). Biallelic TOR1A mutations cause severe arthrogryposis: A case requiring reverse phenotyping. European journal of medical genetics 62, 103544. 10.1016/j.ejmg.2018.09.011.

50. Bressman, S.B., Sabatti, C., Raymond, D., de Leon, D., Klein, C., Kramer, P.L., Brin, M.F., Fahn, S., Breakefield, X., Ozelius, L.J., and Risch, N.J. (2000). The DYT1 phenotype and guidelines for diagnostic testing. Neurology 54, 1746–1752.

51. DeSimone, J.C., Pappas, S.S., Febo, M., Burciu, R.G., Shukla, P., Colon-Perez, L.M., Dauer, W.T., and Vaillancourt, D.E. (2017). Forebrain knock-out of torsinA reduces striatal free-water and impairs whole-brain functional connectivity in a symptomatic mouse model of DYT1 dystonia. Neurobiol Dis 106, 124–132. 10.1016/j.nbd.2017.06.015.

52. Yokoi, F., Dang, M.T., and Li, Y. (2012). Improved motor performance in Dyt1 ΔGAG heterozygous knock-in mice by cerebellar Purkinje-cell specific Dyt1 conditional knocking-out. Behavioural Brain Research 230, 389–398. https://doi.org/10.1016/j.bbr.2012.02.029.

53. Ronzano, R., Lancelin, C., Bhumbra, G.S., Brownstone, R.M., and Beato, M. (2021). Proximal and distal spinal neurons innervating multiple synergist and antagonist motor pools. Elife 10. 10.7554/eLife.70858.

54. Zhang, J., Weinrich, J.A.P., Russ, J.B., Comer, J.D., Bommareddy, P.K., DiCasoli, R.J., Wright, C.V.E., Li, Y., van Roessel, P.J., and Kaltschmidt, J.A. (2017). A Role for Dystonia-Associated Genes in Spinal GABAergic Interneuron Circuitry. Cell reports 21, 666–678. 10.1016/j.celrep.2017.09.079.

55. Liu, Y.B., Tewari, A., Salameh, J., Arystarkhova, E., Hampton, T.G., Brashear, A., Ozelius, L.J., Khodakhah, K., and Sweadner, K.J. (2015). A dystonia-like movement disorder with brain and spinal neuronal defects is caused by mutation of the mouse laminin beta1 subunit, Lamb1. Elife 4. 10.7554/eLife.11102.

56. Nakashima, K., Rothwell, J.C., Day, B.L., Thompson, P.D., Shannon, K., and Marsden, C.D. (1989). Reciprocal inhibition between forearm muscles in patients with writer’s cramp and other occupational cramps, symptomatic hemidystonia and hemiparesis due to stroke. Brain 112, 681–697. 10.1093/brain/112.3.681.

57. Panizza, M., Lelli, S., Nilsson, J., and Hallett, M. (1990). H-reflex recovery curve and reciprocal inhibition of H-reflex in different kinds of dystonia. Neurology 40, 824–828. 10.1212/WNL.40.5.824.

58. Priori, A., Berardelli, A., Mercuri, B., and Manfredi, M. (1995). Physiological effects produced by botulinum toxin treatment of upper limb dystonia. Changes in reciprocal inhibition between forearm muscles. Brain 118 (Pt 3), 801–807.

59. Panizza, M.E., Hallett, M., and Nilsson, J. (1989). Reciprocal inhibition in patients with hand cramps. Neurology 39, 85–89.

60. Brownstone, R.M., and Bui, T.V. (2010). Spinal interneurons providing input to the final common path during locomotion. Prog Brain Res 187, 81–95. 10.1016/b978-0-444-53613-6.00006-x.

61. Krauss, J.K., Yianni, J., Loher, T.J., and Aziz, T.Z. (2004). Deep brain stimulation for dystonia. J Clin Neurophysiol 21, 18–30. 10.1097/00004691-200401000-00004.

62. Herrington, T.M., Cheng, J.J., and Eskandar, E.N. (2016). Mechanisms of deep brain stimulation. J Neurophysiol 115, 19–38. 10.1152/jn.00281.2015.

63. Sherrington, C.S. (1910). Flexion-reflex of the limb, crossed extension-reflex, and reflex stepping and standing. J Physiol 40, 28–121.

64. Maltese, M., Stanic, J., Tassone, A., Sciamanna, G., Ponterio, G., Vanni, V., Martella, G., Imbriani, P., Bonsi, P., Mercuri, N.B., et al. (2018). Early structural and functional plasticity alterations in a susceptibility period of DYT1 dystonia mouse striatum. Elife 7. 10.7554/eLife.33331.

65. Tisch, S., Limousin, P., Rothwell, J.C., Asselman, P., Zrinzo, L., Jahanshahi, M., Bhatia, K.P., and Hariz, M.I. (2006). Changes in forearm reciprocal inhibition following pallidal stimulation for dystonia. Neurology 66, 1091–1093. 10.1212/01.wnl.0000204649.36458.8f.

66. Brownstone, R.M., Bui, T.V., and Stifani, N. (2015). Spinal circuits for motor learning. Current Opinion in Neurobiology 33, 166–173. https://doi.org/10.1016/j.conb.2015.04.007.

67. Forssberg, H., Grillner, S., and Halbertsma, J. (1980). The locomotion of the low spinal cat. I. Coordination within a hindlimb. Acta Physiol Scand 108, 269–281. 10.1111/j.1748-1716.1980.tb06533.x.

68. Forssberg, H., Grillner, S., Halbertsma, J., and Rossignol, S. (1980). The locomotion of the low spinal cat. II. Interlimb coordination. Acta Physiol Scand 108, 283–295. 10.1111/j.1748-1716.1980.tb06534.x.

69. Barbeau, H., and Rossignol, S. (1987). Recovery of locomotion after chronic spinalization in the adult cat. Brain Res 412, 84–95. 10.1016/0006-8993(87)91442-9.

70. Harkema, S., Gerasimenko, Y., Hodes, J., Burdick, J., Angeli, C., Chen, Y., Ferreira, C., Willhite, A., Rejc, E., Grossman, R.G., and Edgerton, V.R. (2011). Effect of epidural stimulation of the lumbosacral spinal cord on voluntary movement, standing, and assisted stepping after motor complete paraplegia: a case study. Lancet 377, 1938–1947. 10.1016/s0140-6736(11)60547-3.

71. Welinder, C., and Ekblad, L. (2011). Coomassie staining as loading control in Western blot analysis. Journal of proteome research 10, 1416–1419. 10.1021/pr1011476.

72. Deerinck, T.J., Bushong, E.A., Lev-Ram, V., Shu, X., Tsien, R.Y., and Ellisman, M.H. (2010). Enhancing Serial Block-Face Scanning Electron Microscopy to Enable High Resolution 3-D Nanohistology of Cells and Tissues. Microscopy and Microanalysis 16, 1138–1139. 10.1017/S1431927610055170.

73. Fox, W.M. (1965). Reflex-ontogeny and behavioural development of the mouse. Animal Behaviour 13, 234–IN235. https://doi.org/10.1016/0003-3472(65)90041-2.

74. Jiang, Z., Carlin, K.P., and Brownstone, R.M. (1999). An in vitro functionally mature mouse spinal cord preparation for the study of spinal motor networks. Brain Res 816, 493–499.

75. Bui, T.V., Akay, T., Loubani, O., Hnasko, T.S., Jessell, T.M., and Brownstone, R.M. (2013). Circuits for grasping: spinal dI3 interneurons mediate cutaneous control of motor behavior. Neuron 78, 191–204. 10.1016/j.neuron.2013.02.007.

76. Özyurt, M.G., Ojeda-Alonso, J., Beato, M., and Nascimento, F. (2022). In vitro longitudinal lumbar spinal cord preparations to study sensory and recurrent motor microcircuits of juvenile mice. 128, 711–726. 10.1152/jn.00184.2022.

77. Dugué, G.P., Dumoulin, A., Triller, A., and Dieudonné, S. (2005). Target-dependent use of co-released inhibitory transmitters at central synapses. J Neurosci 25, 6490–6498. 10.1523/jneurosci.1500-05.2005.

78. Ho, J., Tumkaya, T., Aryal, S., Choi, H., and Claridge-Chang, A. (2019). Moving beyond P values: data analysis with estimation graphics. Nature methods 16, 565–566. 10.1038/s41592-019-0470-3.

79. Pocratsky, A.M., Burke, D.A., Morehouse, J.R., Beare, J.E., Riegler, A.S., Tsoulfas, P., States, G.J.R., Whittemore, S.R., and Magnuson, D.S.K. (2017). Reversible silencing of lumbar spinal interneurons unmasks a task-specific network for securing hindlimb alternation. Nat Commun 8, 1963. 10.1038/s41467-017-02033-x.

80. Pocratsky, A.M., Shepard, C.T., Morehouse, J.R., Burke, D.A., Riegler, A.S., Hardin, J.T., Beare, J.E., Hainline, C., States, G.J.R., Brown, B.L., et al. (2020). Long ascending propriospinal neurons provide flexible, context-specific control of interlimb coordination. eLife 9, e53565. 10.7554/eLife.53565.

