## Supplemental Table 1 for "Pathophysiology of Dyt1 dystonia is mediated by spinal cord dysfunction"

**Title:**

### 7 Table S1

| Figure | Outcome measure<br>Experimental units | Mean ± S.D. (unit) | Normality testing, if applicable | Equality of variance testing, if applicable | Means or ranks comparison(s), where applicable |
|---|---|---|---|---|---|
| Fig.<br>1B<br>Left | **qPCR - *Tor1a***<br>P18 brain samples<br>N = 3 control vs N = 3 cko<br>Experimental unit = # animals | *Con vs cko: (a.u.)*<br>1.00 ± 0.08<br>1.03 ± 0.02 | *Shapiro-Wilk test:*<br>W = 0.99, *P* = 0.78<br>W = 0.75, *P* = 0.13 | *F-test:*<br>F(2,2) = 10.69<br>*P* = 0.17 | *Independent t-test with equal variance:*<br>t(4) = 0.73, *P* = 0.51 |
| Fig.<br>1B<br>Right | **qPCR - *Tor1a***<br>P18 Lumbar spinal cord samples<br>N = 3 control vs N = 3 cko<br>Experimental unit = # animals | *Con vs cko: (a.u.)*<br>1.00 ± 0.04<br>0.04 ± 0.006 | *Shapiro-Wilk test:*<br>W = 0.84, *P* = 0.22<br>W = 0.75, *P* = 0.13 | *F-test:*<br>F(2,2) = 57.00<br>***P* = 0.035** | *Independent t-test with Welch's correction:*<br>t(2.07) = 37.95, ***P* = 0.0006** |
| Fig.<br>1C<br>Right | **Western - torsinA**<br>P18 Lumbar spinal cord samples<br>N = 4 control vs N = 3 cko<br>Experimental unit = # animals | *Con vs cko: (a.u.)*<br>1.13 ± 0.15<br>0.10 ± 0.05 | *Shapiro-Wilk test:*<br>W = 0.90, *P* = 0.44<br>W = 1.00, *P* = 0.93 | *F-test:*<br>F(3,2) = 10.75<br>*P* = 0.17 | *Independent t-test with equal variance:*<br>t(5) = 11.20, ***P* < 0.0001** |
| Fig.<br>2O | **EMG - contractions at rest: total bursting**<br>P17-19<br>N = 4 wildtype control vs N = 6 cko<br>Experimental unit = # animals | *Con vs cko: (%)*<br>0.02 ± 0.03<br>90.17± 5.97 | *Shapiro-Wilk test:*<br>W = 0.93, *P* = 0.55<br>W = 0.75, ***P* = 0.0017** | *F-test:*<br>F(3,5) = 44642<br>***P* < 0.0001** | *Mann-Whitney U test:*<br>U = 0, ***P* = 0.0095** |
| Fig.<br>2P | **EMG - contractions at rest: Ta + G co-activation**<br>P17-19<br>N = 4 wildtype control vs N = 6 cko<br>Experimental unit = # animals | *Con vs cko: (%)*<br>0.52 ± 1.04<br>65.31± 22.83 | *Shapiro-Wilk test:*<br>W = 0.63, ***P* = 0.0012**<br>W = 0.63, ***P* = 0.0009** | *F-test:*<br>F(3,5) = 480.2<br>***P* = 0.0003** | *Mann-Whitney U test:*<br>U = 0, ***P* = 0.0095** |
| Fig.<br>3M | **Fictive locomotion - burst frequency**<br>P1-P5 bilateral caudal<br>N = 5 control vs N = 9 cko<br>Experimental unit = # animals | *Con vs cko: (Hz)*<br>0.25 ± 0.04<br>0.08 ± 0.04 | *Shapiro-Wilk test:*<br>W = 0.94, *P* = 0.64<br>W = 0.81, ***P* = 0.03** | *F-test:*<br>F(10,4) = 3.48<br>*P* = 0.24 | *Mann-Whitney U test:*<br>U = 0, ***P* = 0.001** |
| Fig.<br>3N | **Fictive locomotion - burst frequency**<br>P1-P5 bilateral rostral<br>N = 5 control vs N = 6 cko<br>Experimental unit = # animals | *Con vs cko: (Hz)*<br>0.23 ± 0.04<br>0.07 ± 0.04 | *Shapiro-Wilk test:*<br>W = 0.97, *P* = 0.85<br>W = 0.91, *P* = 0.46 | *F-test:*<br>F(5,4) = 1.046<br>*P* = 0.99 | *Independent t-test with equal variance:*<br>t(9) = 7.26, ***P* < 0.0001** |
| Fig.<br>3O | **Fictive locomotion - burst frequency**<br>P1-P5 ipsilateral rostrocaudal<br>N = 5 control vs N = 11 cko<br>Experimental unit = # animals | *Con vs cko: (Hz)*<br>0.25 ± 0.03<br>0.09 ± 0.05 | *Shapiro-Wilk test:*<br>W = 0.81, *P* = 0.09<br>W = 0.91, *P* = 0.25 | *F-test:*<br>F(10,4) = 2.48<br>*P* = 0.40 | *Independent t-test with equal variance:*<br>t(14) = 6.50, ***P* < 0.0001** |

| Figure | Outcome measure<br>Experimental units | Mean ± S.D. (unit) | Normality testing, if applicable | Equality of variance testing, if applicable | Means or ranks comparison(s), where applicable |
|---|---|---|---|---|---|
| Fig. 3P | **Fictive locomotion - burst coordination**<br>P1-P5 bilateral caudal<br>N = 5 control vs N = 6 cko<br>Experimental unit = # animals | *Con vs cko: (deg) (3-min epochs)*<br>$n_1 = 55$<br>$n_2 = 73$ | *n/a* | *n/a* | *Watson's non-parametric two-sample $U^2$ test:*<br>$U^2 = 2.36$, ***P* < 0.001** |
| Fig. 3Q | **Fictive locomotion - burst coordination**<br>P1-P5 bilateral rostral<br>N = 5 control vs N = 6 cko<br>Experimental unit = # animals | *Con vs cko: (deg) (3-min epochs)*<br>$n_1 = 52$<br>$n_2 = 52$ | *n/a* | *n/a* | *Watson's non-parametric two-sample $U^2$ test:*<br>$U^2 = 1.56$, ***P* < 0.001** |
| Fig. 3R | **Fictive locomotion - burst coordination**<br>P1-P5 ipsilateral rostrocaudal<br>N = 5 control vs N = 11 cko<br>Experimental unit = # animals | *Con vs cko: (deg) (3-min epochs)*<br>$n_1 = 54$<br>$n_2 = 102$ | *n/a* | *n/a* | *Watson's non-parametric two-sample $U^2$ test:*<br>$U^2 = 2.74$, ***P* < 0.001** |

| | | | | | |
|---|---|---|---|---|---|
| Fig. 4D | **Monosynaptic reflex - P7-P8 response duration**<br>Rostral (L1-L3)<br>N = 17 control vs N = 15 cko<br><br>-<br>Caudal (L4-L5)<br>N = 14 control vs N = 14 cko<br>Experimental unit = # roots | *Con vs cko: (ms)*<br><br>*Rostral*<br>2.75 ± 0.66<br>3.61 ± 0.80<br>-<br>*Caudal*<br>2.25 ± 0.45<br>4.06 ± 1.37 | *Shapiro-Wilk test:*<br><br>*Rostral*<br>W = 0.95, *P* = 0.44<br>W = 0.91, *P* = 0.12<br>-<br>*Caudal*<br>W = 0.91, *P* = 0.15<br>W = 0.88, *P* = 0.0502 | *Levene's test: all roots*<br>F(3,56) = 4.11<br>***P* = 0.0104** | *Two-way ANOVA with Geisser-Greenhouse correction: root x genotype*<br>Root: F(1,56) = 0.017, *P* = 0.90<br>Genotype: F(1,56) = 34.64, ***P* < 0.0001**<br>Interaction: F(1,56) = 4.42, ***P* = 0.04**<br>-<br>Tukey's *post hoc* t-test (adjusted *P*):<br>L1-L3 con vs L1-L3 cko: ***P* = 0.038**<br>L4-L5 con vs L4-L5 cko: ***P* < 0.0001**<br>L1-L3 cko vs L4-L5 cko: *P* = 0.52 |
| Fig. 4E | **Monosynaptic reflex - P9-P10 response duration**<br>Rostral (L1-L3)<br>N = 15 control vs N = 19 cko<br><br>-<br>Caudal (L4-L5)<br>N = 11 control vs *N = 12 cko<br>Experimental unit = # roots | *Con vs cko: (ms)*<br><br>*Rostral*<br>2.01 ± 0.43<br>3.26 ± 0.59<br>-<br>*Caudal*<br>1.82 ± 0.18<br>2.78 ± 0.57 | *Shapiro-Wilk test:*<br><br>*Rostral*<br>W = 0.93, *P* = 0.25<br>W = 0.95, *P* = 0.48<br>-<br>*Caudal*<br>W = 0.86, *P* = 0.06<br>W = 0.96, *P* = 0.75 | *Levene's test: all roots*<br>F(3,53) = 2.32<br>*P = 0.085* | *Two-way ANOVA: root x genotype*<br>Root: F(1,53) = 6.21, ***P* = 0.016**<br>Genotype: F(1,53) = 69.90, ***P* < 0.0001**<br>Interaction: F(1,53) = 1.15, *P* = 0.29<br><br>-<br>Tukey's *post hoc* t-test (adjusted *P*):<br>L1-L3 con vs L1-L3 cko: ***P* < 0.0001**<br>L4-L5 con vs L4-L5 cko: ***P* < 0.0001**<br>L1-L3 cko vs L4-L5 cko: *P* = 0.054 |

| Figure | Outcome measure<br>Experimental units | Mean ± S.D. (unit) | Normality testing, if applicable | Equality of variance testing, if applicable | Means or ranks comparison(s), where applicable |
|---|---|---|---|---|---|
| Fig. 4F | **Monosynaptic reflex -**<br>**P11-P12 response duration**<br>Rostral (L1-L3)<br>N = 11 control vs N = 13 cko<br><br>-<br>Caudal (L4-L5)<br>N = 10 control vs N = 9 cko<br>Experimental unit = # roots | *Con vs cko: (ms)*<br><br>*Rostral*<br>1.97 ± 0.94<br>3.74 ± 1.06<br>-<br>*Caudal*<br>1.64 ± 0.41<br>2.97 ± 1.41 | *Shapiro-Wilk test:*<br><br>*Rostral*<br>W = 0.80, ***P* = 0.01**<br>W = 0.83, ***P* = 0.02**<br>-<br>*Caudal*<br>W = 0.95, *P* = 0.70<br>W = 0.92, *P* = 0.36 | *Levene's test: all roots*<br>F(3,39) = 1.52<br>*P = 0.22*<br><br><br>-<br>*F-test: Caudal*<br>F(8,9) = 11.46<br>***P* = 0.0014** | *Mann-Whitney U test: Rostral*<br>U = 14, ***P* = 0.0004**<br><br><br><br>-<br>*Independent t-test with Welch's correction: Caudal*<br>t(2.07) = 37.95, ***P* = 0.0006**<br>-<br>*Mann-Whitney U test: Rostral vs Caudal*<br>U = 31, ***P* = 0.071** |
| Fig. 4G | **Monosynaptic reflex -**<br>**P7-P8 latency**<br>Rostral (L1-L3)<br>N = 16 control vs N = 15 cko<br><br>-<br>Caudal (L4-L5)<br>N = 14 control vs N = 14 cko<br>Experimental unit = # roots | *Con vs cko: (ms)*<br><br>*Rostral*<br>4.50 ± 0.46<br>6.83 ± 0.78<br>-<br>*Caudal*<br>4.69 ± 0.81<br>7.55 ± 1.41 | *Shapiro-Wilk test:*<br><br>*Rostral*<br>W = 0.92, *P* = 0.14<br>W = 0.95, *P* = 0.60<br>-<br>*Caudal*<br>W = 0.90, *P* = 0.13<br>W = 0.97, *P* = 0.83 | *Levene's test: all roots*<br>F(3,55) = 3.45<br>***P* = 0.023** | *Two-way ANOVA with Geisser-Greenhouse correction: root x genotype*<br>Root: F(1,56) = 4.04, ***P* = 0.049**<br>Genotype: F(1,56) = 123.2, ***P* < 0.0001**<br>Interaction: F(1,56) = 1.15, *P* = 0.29<br>-<br>Tukey's *post hoc* t-test (adjusted *P*):<br>L1-L3 con vs L1-L3 cko: ***P* < 0.0001**<br>L4-L5 con vs L4-L5 cko: ***P* < 0.0001**<br>L1-L3 cko vs L4-L5 cko: *P* = 0.15 |
| Fig. 4G | **Monosynaptic reflex -**<br>**P9-P10 latency**<br>Rostral (L1-L3)<br>N = 15 control vs N = 19 cko<br><br>-<br>Caudal (L4-L5)<br>N = 11 control vs N = 14 cko<br>Experimental unit = # roots | *Con vs cko: (ms)*<br>*Rostral*<br>3.87 ± 0.41<br>5.28 ± 0.60<br><br>-<br>*Caudal*<br>3.81 ± 0.44<br>6.09 ± 0.95 | *Shapiro-Wilk test:*<br>*Rostral*<br>W = 0.96, *P* = 0.64<br>W = 0.96, *P* = 0.51<br><br>-<br>*Caudal*<br>W = 0.94, *P* = 0.50<br>W = 0.92, *P* = 0.20 | *Levene's test: all roots*<br>F(3,55) = 4.25<br>***P* = 0.009** | *Two-way ANOVA with Geisser-Greenhouse correction: root x genotype*<br>Root: F(1,55) = 4.81, ***P* = 0.033**<br>Genotype: F(1,55) = 118.1, ***P* < 0.0001**<br>Interaction: F(1,55) = 6.41, ***P* = 0.014**<br>-<br>Tukey's *post hoc* t-test (adjusted *P*):<br>L1-L3 con vs L1-L3 cko: ***P* < 0.0001**<br>L4-L5 con vs L4-L5 cko: ***P* < 0.0001**<br>L1-L3 cko vs L4-L5 cko: ***P* = 0.0042** |

| Figure | Outcome measure<br>Experimental units | Mean ± S.D. (unit) | Normality testing, if applicable | Equality of variance testing, if applicable | Means or ranks comparison(s), where applicable |
|---|---|---|---|---|---|
| Fig. 4G | **Monosynaptic reflex - P11-P12 latency**<br>Rostral (L1-L3)<br>N = 11 control vs N = 13 cko<br><br>-<br>Caudal (L4-L5)<br>N = 10 control vs N = 9 cko<br>Experimental unit = # roots | *Con vs cko: (ms)*<br>*Rostral*<br>3.16 ± 0.23<br>5.54 ± 0.80<br><br>-<br>*Caudal*<br>3.34 ± 0.30<br>6.04 ± 0.95 | *Shapiro-Wilk test:*<br>*Rostral*<br>W = 0.95, *P* = 0.60<br>W = 0.96, *P* = 0.68<br><br>-<br>*Caudal*<br>W = 0.94, *P* = 0.52<br>W = 0.91, *P* = 0.29 | *Levene's test: all roots*<br>F(3,39) = 5.96<br>***P* = 0.002** | *Two-way ANOVA with Geisser-Greenhouse correction: root x genotype*<br>Root: F(1,39) = 3.01, ***P* = 0.09**<br>Genotype: F(1,39) = 164.1, ***P* < 0.0001**<br>Interaction: F(1,39) = 0.68, *P* = 0.41<br>-<br>Tukey's *post hoc* t-test (adjusted *P*):<br>L1-L3 con vs L1-L3 cko: ***P* < 0.0001**<br>L4-L5 con vs L4-L5 cko: ***P* < 0.0001**<br>L1-L3 cko vs L4-L5 cko: *P* = 0.28 |

| Figure | Outcome measure<br>Experimental units | Mean ± S.D. (unit) | Normality testing, if applicable | Equality of variance testing, if applicable | Means or ranks comparison(s), where applicable |
|---|---|---|---|---|---|
| Fig. 5C | **Intrinsic Mn properties - P1-P13 Resting membrane potential**<br>N = 52 control vs N = 77 cko<br>Experimental unit = # Mns | *Con vs cko: (mV)*<br>-63.40 ± 5.20<br>-63.47 ± 4.60 | *Shapiro-Wilk test:*<br>W = 0.95, ***P* = 0.019**<br>W = 0.97, *P* = 0.13 | *F-Test:*<br>F(51,76) = 1.28<br>*P* = 0.32 | *Mann-Whitney U test:*<br>U = 2089, *P* = 0.67 |
| Fig. 5D | **Intrinsic Mn properties - P1-P13 Whole cell capacitance**<br>N = 73 control vs N = 108 cko<br>Experimental unit = # Mns | *Con vs cko: (pF)*<br>309.96 ± 107.49<br>146.98 ± 52.74 | *Shapiro-Wilk test:*<br>W = 0.97, *P* = 0.08<br>W = 0.93, ***P* < 0.0001** | *F-Test:*<br>F(72,107) = 4.15<br>***P* < 0.0001** | *Mann-Whitney U test:*<br>U = 7350, ***P* < 0.0001** |
| Fig. 5E | **Intrinsic Mn properties - P1-P13 Input resistance**<br>N = 73 control vs N = 111 cko<br>Experimental unit = # Mns | *Con vs cko: (MΩ)*<br>39.13 ± 25.00<br>178.09 ± 111.9 | *Shapiro-Wilk test:*<br>W = 0.81, ***P* < 0.0001**<br>W = 0.89, ***P* < 0.0001** | *F-Test:*<br>F(72,110) = 4.15<br>***P* < 0.0001** | *Mann-Whitney U test:*<br>U = 343, ***P* < 0.0001** |
| Fig. 5G | **DR-evoked EPSC - P1-P13 Area**<br>N = 61 control vs N = 130 cko<br>Experimental unit = # root responses | *Con vs cko: (pA.ms)*<br>2.05e4 ± 1.89e4<br>3.99e3 ± 5.41e3 | *Shapiro-Wilk test:*<br>W = 0.84, ***P* < 0.0001**<br>W = 0.70, ***P* < 0.0001** | *F-Test:*<br>F(60,129) = 12.14<br>***P* < 0.0001** | *Mann-Whitney U test:*<br>U = 6942, ***P* < 0.0001** |
| Fig. 5H | **DR-evoked EPSC - P1-P13 Conductance**<br>N = 61 control vs N = 128 cko<br>Experimental unit = # root responses | *Con vs cko: (nS)*<br>18.94 ± 16.97<br>4.25 ± 4.22 | *Shapiro-Wilk test:*<br>W = 0.79, ***P* < 0.0001**<br>W = 0.76, ***P* < 0.0001** | *F-Test:*<br>F(60,127) = 16.19<br>***P* < 0.0001** | *Mann-Whitney U test:*<br>U = 6955, ***P* < 0.0001** |
| Fig. 5I | **DR-evoked EPSC - P1-P13 Conductance scaled**<br>N = 61 control vs N = 126 cko<br>Experimental unit = # root responses | *Con vs cko: (%)*<br>79.50 ± 78.40<br>68.52 ± 91.98 | *Shapiro-Wilk test:*<br>W = 0.71, ***P* < 0.0001**<br>W = 0.55, ***P* < 0.0001** | *F-Test:*<br>F(60,125) = 0.73<br>*P* = 0.17 | *Mann-Whitney U test:*<br>U = 4499, *P* = 0.059 |
| Fig. 5J | **DR-evoked EPSC - P1-P13 Latency**<br>N = 60 control vs N = 131 cko<br>Experimental unit = # root responses | *Con vs cko: (ms)*<br>6.06 ± 3.92<br>11.51 ± 6.74 | *Shapiro-Wilk test:*<br>W = 0.81, ***P* < 0.0001**<br>W = 0.91, ***P* < 0.0001** | *F-Test:*<br>F(59,130) = 0.34<br>***P* < 0.0001** | *Mann-Whitney U test:*<br>U = 1628, ***P* < 0.0001** |

| Figure | Outcome measure<br>Experimental units | Mean ± S.D. (unit) | Normality testing, if applicable | Equality of variance testing, if applicable | Means or ranks comparison(s), where applicable |
|---|---|---|---|---|---|
| Fig. 5M | **L4 & L5 DR conduction velocity**<br>P6-P10<br>N = 15 control vs N = 15 cko<br>Experimental unit = # dorsal roots | *Con vs cko: (m/s)*<br>4.73 ± 1.70<br>1.83 ± 0.76 | *Shapiro-Wilk test:*<br>W = 0.97, *P* = 0.89<br>W = 0.96, *P* = 0.63 | *F-Test:*<br>F(14,14) = 5.06<br>***P* = 0.0045** | *Independent t-test with Welch's correction:*<br>t(19.3) = 6.03, ***P* < 0.0001** |
| Fig. 5P | **L4 & L5 DR microstim. scaled conduction time - average**<br>P6-P10<br>[#]N = 14 control vs N = 15 cko<br>Experimental unit = # dorsal roots | *Con vs cko: (ms/mm)*<br>0.38 ± 0.11<br>0.89 ± 0.25 | *Shapiro-Wilk test:*<br>W = 0.95, *P* = 0.63<br>W = 0.88, ***P* = 0.048** | *F-test:*<br>F(13,14) = 5.31<br>***P* = 0.0047** | *Mann-Whitney U test:*<br>U = 0, ***P* < 0.0001** |
| Fig. 5P | **L4 & L5 DR microstim. scaled conduction time - stand. deviation**<br>P6-P10<br>[#]N = 14 control vs N = 15 cko<br>Experimental unit = # dorsal roots | *Con vs cko: (ms/mm)*<br>0.11 ± 0.06<br>0.30 ± 0.14 | *Shapiro-Wilk test:*<br>W = 0.91, *P* = 0.13<br>W = 0.85, ***P* = 0.019** | *F-test:*<br>F(13,14) = 4.40<br>***P* = 0.0112** | *Mann-Whitney U test:*<br>U = 12, ***P* < 0.0001** |

| | | | | | |
|---|---|---|---|---|---|
| Fig. S1D | **qPCR - *Tor1a***<br>P18 DRG samples<br>N = 4 control vs N = 4 cko<br>Experimental unit = # animals | *Con vs cko: (a.u.)*<br>1.00± 0.08<br>0.03 ± 0.008 | *Shapiro-Wilk test:*<br>W = 0.97, *P* = 0.85<br>W = 0.94, *P* = 0.68 | *F-test:*<br>F(3,3) = 85.00<br>***P* = 0.0042** | *Independent t-test with Welch's correction:*<br>t(3.071) = 25.62, ***P* = 0.0001** |
| Fig. S1E | **qPCR - *Tor1a***<br>P18 Heart samples<br>N = 4 control vs N = 4 cko<br>Experimental unit = # animals | *Con vs cko: (a.u.)*<br>1.03± 0.28<br>1.18 ± 0.07 | *Shapiro-Wilk test:*<br>W = 0.88, *P* = 0.36<br>W = 0.88, *P* = 0.33 | *F-test:*<br>F(3,3) = 16.43<br>***P* = 0.046** | *Independent t-test with Welch's correction:*<br>t(3.364) = 1.101, *P* = 0.34 |
| Fig. S1F | **qPCR - *Tor1a***<br>P18 Liver samples<br>N = 4 control vs N = 4 cko<br>Experimental unit = # animals | *Con vs cko: (a.u.)*<br>1.04± 0.32<br>0.96 ± 0.07 | *Shapiro-Wilk test:*<br>W = 0.92, *P* = 0.54<br>W = 0.95, *P* = 0.74 | *F-test:*<br>F(3,3) = 21.59<br>***P* = 0.0312** | *Independent t-test with Welch's correction:*<br>t(3.277) = 0.48, *P* = 0.66 |

| Figure | Outcome measure<br>Experimental units | Mean ± S.D. (unit) | Normality testing, if applicable | Equality of variance testing, if applicable | Means or ranks comparison(s), where applicable |
|---|---|---|---|---|---|
| Fig. S1G | **Postnatal body weight**<br>P1-P14<br>N = 8 control vs N = 6 cko<br>Experimental unit = # animals | *Con vs cko: (g)*<br>*P1-P2*<br>2.22 ± 0.23<br>1.86 ± 0.37<br>-<br>*P3-P4*<br>3.13 ± 0.64<br>2.50 ± 0.55<br>-<br>*P5-P6*<br>4.00 ± 0.54<br>3.00 ± 0.00<br>-<br>*P7-P8*<br>5.13 ± 0.64<br>3.50 ± 0.55<br>-<br>*P9-P10*<br>6.50 ± 0.54<br>3.67 ± 0.52<br>-<br>*P11-P12*<br>7.63 ± 0.74<br>4.17 ± 0.98<br>-<br>*P13-P14*<br>8.63 ± 0.74<br>4.50 ± 0.84 | *Shapiro-Wilk test:*<br>*P1-P2*<br>W = 0.89, *P* = 0.25<br>W = 0.84, *P* = 0.14<br>-<br>*P3-P4*<br>W = 0.81, ***P* = 0.04**<br>W = 0.68, ***P* = 0.004**<br>-<br>*P5-P6*<br>W = 0.73, ***P* = 0.0052**<br>W = 0.82, *P* = 0.09<br>-<br>*P7-P8*<br>W = 0.81, ***P* = 0.037**<br>W = 0.68, ***P* = 0.004**<br>-<br>*P9-P10*<br>W = 0.66, ***P* = 0.0009**<br>W = 0.64, ***P* = 0.0014**<br>-<br>*P11-P12*<br>W = 0.80, ***P* = 0.027**<br>W = 0.78, ***P* = 0.035**<br>-<br>*P13-P14*<br>W = 0.80, ***P* = 0.027**<br>W = 0.70, ***P* = 0.006** | *F-Test:*<br>*P1-P2*<br>F(7,5) = 2.67<br>*P* = 0.23<br>-<br>*P3-P4*<br>F(7,5) = 1.37<br>*P* = 0.75<br>-<br>*P5-P6*<br>F(7,5) = ∞<br>***P* < 0.0001**<br>-<br>*P7-P8*<br>F(7,5) = 1.37<br>*P* = 0.75<br>-<br>*P9-P10*<br>F(7,5) = 1.07<br>*P* = 0.97<br>-<br>*P11-P12*<br>F(7,5) = 1.75<br>*P* = 0.49<br>-<br>*P13-P14*<br>F(7,5) = 1.27<br>*P* = 0.75 | *Mann-Whitney U test:*<br>*P1-P2*<br>U = 11, *P* = 0.11<br><br>-<br>*P3-P4*<br>U = 11, *P* = 0.14<br><br>-<br>*P5-P6*<br>U = 3, ***P* = 0.0047**<br><br>-<br>*P7-P8*<br>U = 1.5, ***P* = 0.0013**<br><br>-<br>*P9-P10*<br>U = 0, ***P* = 0.0007**<br><br>-<br>*P11-P12*<br>U = 0, ***P* = 0.0007**<br><br>-<br>*P13-P14*<br>U = 0, ***P* = 0.0007** |
| Fig. S1X | **Ultrastructure - affected DRG neurons**<br>P18 L5 DRG<br>N = 4 control vs N = 4 cko<br>Experimental unit = # animals | *Con vs cko: (% total)*<br>0.00 ± 0.00<br>8.03 ± 4.96 | *Shapiro-Wilk test:*<br>W = n/a, *P* = n/a<br>W = 0.77, *P* = 0.06 | *n/a* | *Mann-Whitney U test:*<br>U = 0, ***P* = 0.03** |
| Fig. S1Y | **Ultrastructure - affected spinal neurons**<br>P18 L5 spinal cord<br>N = 4 control vs N = 4 cko<br>Experimental unit = # animals | *Con vs cko: (% total)*<br>0.00 ± 0.00<br>59.12 ± 5.76 | *Shapiro-Wilk test:*<br>W = n/a, *P* = n/a<br>W = 0.93, *P* = 0.57 | *n/a* | *Mann-Whitney U test:*<br>U = 0, ***P* = 0.03** |

| Figure | Outcome measure<br>Experimental units | Mean ± S.D. (unit) | Normality testing, if applicable | Equality of variance testing, if applicable | Means or ranks comparison(s), where applicable |
|---|---|---|---|---|---|
| Fig. S3B | **Fictive locomotion - burst frequency**<br>P1-P5 unilateral rostral<br>N = 5 control vs N = 4 cko<br>Experimental unit = # animals | *Con vs cko: (Hz)*<br>0.24 ± 0.02<br>0.08 ± 0.04 | *Shapiro-Wilk test:*<br>W = 0.96, *P* = 0.82<br>W = 0.96, *P* = 0.79 | *F-Test:*<br>F(3,4) = 4.89<br>*P* = 0.16 | *Independent t-test with equal variance:*<br>t(7) = 8.09, ***P* < 0.0001** |
| Fig. S3C | **Fictive locomotion - burst frequency**<br>P1-P5 unilateral caudal<br>N = 5 control vs N = 11 cko<br>Experimental unit = # animals | *Con vs cko: (Hz)*<br>0.24 ± 0.08<br>0.04 ± 0.04 | *Shapiro-Wilk test:*<br>W = 0.94, *P* = 0.67<br>W = 0.87, *P* = 0.08 | *F-test:*<br>F(10,4) = 1.24<br>*P* = 0.90 | *Independent t-test with equal variance:*<br>t(14) = 7.72, ***P* < 0.0001** |
| Fig. S3D | **Fictive locomotion - burst cycle duration**<br>P1-P5 unilateral rostral<br>N = 5 control vs N = 11 cko<br>Experimental unit = # animals | *Con vs cko: (s)*<br>4.64 ± 0.17<br>16.44 ± 10.28 | *Shapiro-Wilk test:*<br>W = 0.80, *P* = 0.08<br>W = 0.89, *P* = 0.15 | *F-test:*<br>F(10,4) = 3577<br>***P* < 0.0001** | *Independent t-test with Welch's correction:*<br>t(10.01) = 3.81, ***P* = 0.0034** |
| Fig. S3E | **Fictive locomotion - burst cycle duration**<br>P1-P5 unilateral caudal<br>N = 5 control vs N = 11 cko<br>Experimental unit = # animals | *Con vs cko: (s)*<br>4.26 ± 0.64<br>18.01 ± 8.70 | *Shapiro-Wilk test:*<br>W = 0.97, *P* = 0.85<br>W = 0.88, *P* = 0.10 | *F-test:*<br>F(10,4) = 186.6<br>***P* = 0.0001** | *Independent t-test with Welch's correction:*<br>t(10.23) = 5.21, ***P* = 0.0004** |
| Fig. S3F | **Fictive locomotion - burst cycle duration**<br>P1-P5 bilateral caudal<br>N = 5 control vs N = 9 cko<br>Experimental unit = # animals | *Con vs cko: (s)*<br>4.20 ± 0.57<br>17.77 ± 7.65 | *Shapiro-Wilk test:*<br>W = 0.94, *P* = 0.66<br>W = 0.95, *P* = 0.70 | *F-test:*<br>F(8,4) = 178.5<br>***P* = 0.0002** | *Independent t-test with Welch's correction:*<br>t(8.16) = 5.29, ***P* = 0.0007** |
| Fig. S3G | **Fictive locomotion - burst cycle duration**<br>P1-P5 bilateral rostral<br>N = 5 control vs N = 6 cko<br>Experimental unit = # animals | *Con vs cko: (s)*<br>4.51 ± 0.84<br>21.25 ± 10.21 | *Shapiro-Wilk test:*<br>W = 0.97, *P* = 0.88<br>W = 0.96, *P* = 0.83 | *F-test:*<br>F(5,4) = 149.4<br>***P* = 0.0002** | *Independent t-test with Welch's correction:*<br>t(5.08) = 4.0, ***P* = 0.01** |
| Fig. S3H | **Fictive locomotion - burst cycle duration**<br>P1-P5 ipsilateral rostrocaudal<br>N = 5 control vs N = 11 cko<br>Experimental unit = # animals | *Con vs cko: (s)*<br>4.06 ± 0.54<br>17.00 ± 10.35 | *Shapiro-Wilk test:*<br>W = 0.81, *P* = 0.10<br>W = 0.89, *P* = 0.12 | *F-test:*<br>F(10,4) = 365.8<br>***P* < 0.0001** | *Independent t-test with Welch's correction:*<br>t(10.12) = 4.14, ***P* = 0.002** |
| Fig. S3I | **Fictive locomotion - cross-coherence**<br>P1-P5 bilateral caudal<br>N = 5 control vs N = 9 cko<br>Experimental unit = # animals | *Con vs cko: (a.u.)*<br>0.88 ± 0.05<br>0.84 ± 0.03 | *Shapiro-Wilk test:*<br>W = 0.90, *P* = 0.39<br>W = 0.91, *P* = 0.29 | *F-test:*<br>F(4,8) = 1.842<br>*P* = 0.43 | *Independent t-test with equal variance:*<br>t(12) = 1.85, ***P* = 0.09** |

| Figure | Outcome measure<br>Experimental units | Mean ± S.D. (unit) | Normality testing, if applicable | Equality of variance testing, if applicable | Means or ranks comparison(s), where applicable |
|---|---|---|---|---|---|
| Fig. S3J | **Fictive locomotion - cross-coherence**<br>P1-P5 bilateral rostral<br>N = 5 control vs N = 6 cko<br>Experimental unit = # animals | *Con vs cko: (Hz)*<br>0.90 ± 0.05<br>0.84 ± 0.02 | *Shapiro-Wilk test:*<br>W = 0.91, *P* = 0.44<br>W = 0.97, *P* = 0.92 | *F-test:*<br>F(4,5) = 3.78<br>*P* = 0.18 | *Independent t-test with equal variance:*<br>t(9) = 2.48, ***P* = 0.04** |
| Fig. S6K | **Fictive locomotion - cross-coherence**<br>P1-P5 ipsilateral rostrocaudal<br>N = 5 control vs N = 11 cko<br>Experimental unit = # animals | *Con vs cko: (Hz)*<br>0.89 ± 0.03<br>0.83 ± 0.03 | *Shapiro-Wilk test:*<br>W = 0.89, *P* = 0.33<br>W = 0.89, *P* = 0.16 | *F-test:*<br>F(10,4) = 1.37<br>*P* = 0.82 | *Independent t-test with equal variance:*<br>t(14) = 4.22, ***P* = 0.0009** |

| | | | | | |
|---|---|---|---|---|---|
| Fig. S5A | **DR-evoked EPSC area vs age - slope for line of best fit**<br>P1-P13<br>N = 60 control vs N = 131 cko<br>Experimental unit = # responses | *Con vs cko: (slope)*<br>Y = -1.60e3x + 2.88e4; $R^2$ = 0.0660<br>Y = -6.85e2x + 9.34e3; $R^2$ = 0.202 | *n/a* | *n/a* | *Fisher's r to Z test:*<br>Z = 1.39, t = 9.2, ***P* < 0.001** |
| Fig. S5A | **DR-evoked EPSC latency vs age - slope for line of best fit**<br>P1-P13<br>N = 60 control vs N = 131 cko<br>Experimental unit = # responses | *Con vs cko: (slope)*<br>Y = -0.71x + 9.58; $R^2$ = 0.242<br>Y = -0.63x + 16.47; $R^2$ = 0.108 | *n/a* | *n/a* | *Fisher's r to Z test:*<br>Z = 1.24, t = 5.8, ***P* < 0.001** |
| Fig. S5G | **L4 & L5 VR whole root conduction velocity**<br>P6-P9<br>N = 10 control vs N = 10 cko<br>Experimental unit = # roots | *Con vs cko: (m/s)*<br>4.91 ± 1.97<br>3.09 ± 1.51 | *Shapiro-Wilk test:*<br>W = 0.91, *P* = 0.28<br>W = 0.86, *P* = 0.07 | *F-test:*<br>F(9,9) = 1.69<br>*P* = 0.44 | *Independent t-test with equal variance:*<br>t(18) = 2.31, ***P* = 0.033** |
| Fig. S5I | **L4 & L5 VR microstim. conduction time - average**<br>P6-P9<br>N = 10 control vs #N = 9 cko | *Con vs cko: (ms/mm)*<br>0.42 ± 0.20<br>0.71 ± 0.25 | *Shapiro-Wilk test:*<br>W = 0.77, ***P* = 0.007**<br>W = 0.87, *P* = 0.13 | *F-test:*<br>F(9,8) = 1.58<br>*P* = 0.51 | *Mann-Whitney U test:*<br>U = 13, ***P* = 0.0076** |
| Fig. S5I | **L4 & L5 VR microstim. conduction time - standard deviation**<br>P6-P9<br>#N = 10 control vs #N = 9 cko | *Con vs cko: (ms/mm)*<br>0.13 ± 0.13<br>0.31 ± 0.19 | *Shapiro-Wilk test:*<br>W = 0.75, ***P* = 0.004**<br>W = 0.86, *P* = 0.103 | *F-test:*<br>F(9,8) = 2.072<br>*P* = 0.30 | *Mann-Whitney U test:*<br>U = 18, ***P* = 0.028** |

| Figure | Outcome measure<br>Experimental units | Mean ± S.D. (unit) | Normality testing,<br>if applicable | Equality of variance testing, if applicable | Means or ranks comparison(s), where applicable |
|---|---|---|---|---|---|
| Movie S7 | **qPCR - *Tor1a***<br>P18 spinal samples<br>N = 3 control vs N = 3 DRG cko<br>Experimental unit = # animals | *Con vs cko: (a.u.)*<br>1.22 ± 0.04<br>1.21 ± 0.03 | *Shapiro-Wilk test:*<br>W = 0.99, *P* = 0.82<br>W = 0.86, *P* = 0.28 | *F-Test:*<br>F(2,2) = 1.68<br>*P* = 0.75 | *Independent t-test with equal variance:*<br>t(4) = 0.115, *P* = 0.91 |
| Movie S7 | **qPCR - *Tor1a***<br>P18 DRG samples<br>N = 3 control vs N = 3 DRG cko<br>Experimental unit = # animals | *Con vs cko: (a.u.)*<br>1.17 ± 0.02<br>0.19 ± 0.008 | *Shapiro-Wilk test:*<br>W = 0.99, *P* = 0.83<br>W = 0.94, *P* = 0.54 | *F-Test:*<br>F(2,2) = 7.63<br>*P* = 0.23 | *Independent t-test with equal variance:*<br>t(4) = 69.77, ***P* < 0.0001** |
